# Seven gut microbiome types across two Japanese cohorts defined by co-abundance-based microbial guilds and associated with host physiology

**DOI:** 10.64898/2026.09.07.749831

**Authors:** Jiro Nakayama, Misako Yoden, Ayaka Uchikawa, Masaru Tanaka, Shiori Tamura, Akari Shinoda, Rie Momoda, Keishi Kameyama, Takako Inoue, Hisayoshi Watanabe, Yasuto Tanaka

## Abstract

Intensive efforts have been made to capture structural features of the gut microbiome for assessing microbiota status in relation to host health. However, gut microbiome function emerges not only from individual microbial taxa but also from their organization into ecological guilds, complicating microbiome structure-function relationships. To address this challenge, we applied a co-abundance-based analytical framework to identify gut microbiome types (GMTs) across a discovery cohort (D-cohort) and an independent validation cohort (V-cohort) of Japanese adults. Using paired fecal samples collected one month apart from 116 participants in the D-cohort, we identified 27 co-abundance groups (CAGs) based on 16S rRNA amplicon sequence variants. These CAGs defined seven GMTs (MT-1 to MT-7) with characteristic short-chain fatty acid and bile acid profiles. The GMTs showed temporal stability, with more than 70% of participants remaining in the same GMT across sampling points. The 27-CAG framework classified 404 participants in the independent V-cohort into the seven GMTs, which represented reproducible microbial ecological states across cohorts. MT-4 exhibited dysbiosis-like features, including reduced microbial diversity, depletion of *Faecalibacterium* and butyrate, and enrichment of inflammation-associated taxa, and was associated with higher alcohol consumption and elevated γ-glutamyl transferase levels. In contrast, *Bifidobacterium*-enriched MT-5 was associated with favorable renal function profiles. Collectively, co-abundance-defined microbial guilds capture coherent ecological and functional states of the gut microbiota, providing a transferable framework for linking gut microbiome organization to host physiology beyond conventional taxonomy-based classification.

## 1. Introduction

The highly diverse structure of the human gut microbiota has been extensively analyzed, largely based on sequence variation in the 16S rRNA gene (Abdill et al., 2025). In particular, QIIME2, a bioinformatics platform used with curated reference databases such as SILVA, enables profiling of microbial communities at the amplicon sequence variant (ASV) level (Bolyen et al., 2019a; Wang et al., 2025). These approaches enable high-resolution and standardized taxonomic profiling across cohorts, facilitating large-scale cataloging of the gut microbiome in relation to host phenotypes and genotypes (Falony et al., 2016; Rothschild et al., 2018). Such taxonomic frameworks have become a cornerstone for exploring associations between the gut microbiome and host health and disease (Falony et al., 2016; Li et al., 2025; Rothschild et al., 2018; Su et al., 2022).

In addition, clustering approaches have been widely used to classify variation in the human gut microbiota across individuals. Notably, the concept of enterotypes provided a framework for global gut microbiome typing (MetaHIT Consortium (additional members) et al., 2011). Subsequently, many studies have applied the enterotype framework to characterize inter-individual variation in the gut microbiota within or across populations and to explore its associations with host health and disease Christensen et al., 2020; Takagi et al., 2022).

However, taxonomic composition alone provides only a partial view of microbiome function because community behavior and host-related functions can emerge from ecological interactions among microorganisms (Culp and Goodman, 2023; Faust and Raes, 2012). These interactions, including metabolic cross-feeding, niche partitioning, and competitive or cooperative relationships, collectively shape the ecological organization of the gut microbiome. Thus, understanding the microbiome as an ecological system is important for linking microbial community structure to its functional and physiological relevance to the host.

To address this challenge, co-abundance-based approaches provide an effective strategy to capture ecological relationships beyond taxonomy (Wu et al., 2021). Guild-based analyses have successfully identified coordinated microbial groups associated with predefined health and disease phenotypes, culminating in the identification of two competing guilds that showed stable relationships across diverse populations and disease states (Wu et al., 2024). Phenotype-independent approaches have also revealed generalizable microbial guilds, such as enterosignatures, that describe gut microbiome variation as combinations of co-occurring bacterial groups (Frioux et al., 2023). Nevertheless, integrating multiple co-abundance guilds to characterize gut microbiome configurations and their associations with varying health status among individuals in the general population remains an important area of investigation.

Here, we present a co-abundance-based analytical framework that integrates ASV-level co-abundance groups (CAGs) into gut microbiome types (GMTs) and links these community states to microbial metabolites and host physiological characteristics. Using paired fecal samples from a discovery cohort (D-cohort), we identified 27 CAGs that collectively resolved seven GMTs, representing coordinated microbial ecosystem states. We then applied the 27-CAG and 7-GMT framework to the independent validation cohort (V-cohort) without relearning to assess the transferability of the identified ecological and host-associated characteristics across cohorts.

## 2. Materials and Methods

### 2.1. Study design and cohorts

This study was conducted using two independent Japanese cohorts: a discovery cohort (D-cohort) and an external validation cohort (V-cohort). To capture natural inter-individual variation in the Japanese gut microbiome, the cohorts were designed with limited exclusion criteria and included community-dwelling subjects with diverse physiological and clinical backgrounds.

In the D-cohort, participants were instructed to defer stool collection if they had experienced influenza or acute gastroenteritis within the preceding two weeks or were currently symptomatic. In cases of diarrhea, stool samples were collected after symptoms had resolved. D-cohort included 116 community-dwelling adults aged 19-76 years in Japan (Table 1). Fecal samples were collected twice at a one-month interval (T0 and T1) and analyzed for gut microbiome composition, short-chain fatty acids (SCFAs), and bile acids (BAs). Dietary intake, including alcohol consumption, was assessed using a validated self-administered diet history questionnaire (DHQ) (Kobayashi et al., 2012, 2011) and additional information on medical history, current medication use, smoking habits, and bowel conditions was collected via questionnaires.

**Table 1.** Characteristics of study participants.

| Variable | D-cohort | V-cohort |
| --- | --- | --- |
| No. of subjects | 116 | 404 |
| No. of samples<sup>1</sup> | 232 | 404 |
| Age, years (mean ± std [range]) | 41.9 ± 14.3 (19 – 78) | 53.1 ± 5.9 (34 – 65) |
| Sex, n (male / female) | 72 / 44 | 289 / 115 |
| BMI (kg/m<sup>2</sup>) | 22.8 ± 3.7 | 25.7 ± 3.7 |
| Current smoker, n (%) | 9 (7.8%) | 58 (14.4%) |
| Current drinker, n (%)<br>(low / moderate / high)<sup>2</sup> | 36 (31.6%) / 53 (46.5%) / 25 (21.9%)<sup>3</sup> | 116 (28.7) / 174 (43.0) / 114 (28.2%) |

<sup>1</sup>In D-cohort, samples were collected from each participant at two time points with a one-month interval.
<sup>2</sup>In D-discovery cohort, alcohol consumption categories were defined based on estimated intake (<1, 1–20, and >20 g/day) derived from the DHQL questionnaire. In V-cohort, categories were defined based on self-reported drinking frequency (“almost none”, “occasionally”, and “almost daily”).
<sup>3</sup>Data for three participants were missing due to non-response in the DHQL questionnaire.

V-cohort consisted of 404 community-dwelling adults aged 42-62 years undergoing routine health check-ups. Fecal samples were collected once and analyzed for microbiome composition, SCFAs, and BAs. Clinical measurements obtained during the health check-ups were used to evaluate association analyses between the gut microbiome and host clinical and physiological characteristics. Values outside the reference range for clinical parameters were defined according to standard reference ranges used in routine health check-ups in Japan (Supplementary Table S12). The workflow of this study is shown in Figure 1. Detailed sampling procedures and participant characteristics are described in the Supplementary Note 1 and Supplementary Tables S1, S2, and S11, respectively.

**Figure 1.**
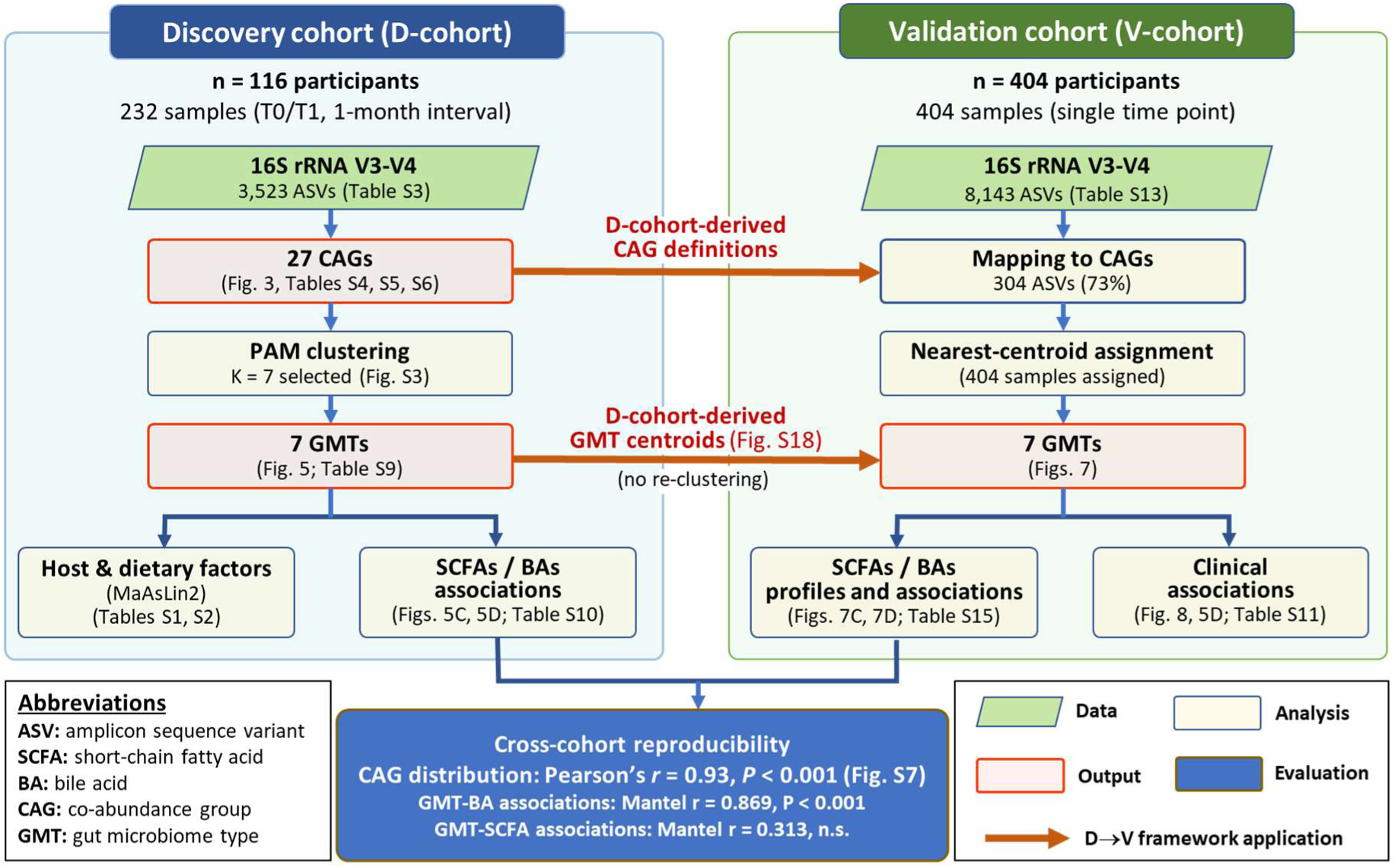
Study design and analytical workflow for co-abundance group (CAG)-based gut microbiome typing. In the D-cohort, fecal samples were collected longitudinally at two time points separated by a one-month interval (T0 and T1) from 116 Japanese adults and subjected to 16S rRNA amplicon sequencing and co-abundance network analysis to define 27 co-abundance groups (CAGs) at the ASV level. Based on CAG abundance profiles, samples were classified into seven gut microbiome types (GMTs), which were subsequently analyzed in relation to microbial metabolites, dietary intake, and host physiological parameters. In the independent validation cohort, ASVs were mapped to the 27 CAGs identified in the discovery cohort, and samples were classified to the seven GMTs to evaluate reproducibility and associations with clinical parameters. Corresponding figures and supplementary tables are indicated in parentheses.

### 2.2. Fecal sample collection

For bacterial composition analysis, fecal samples were collected from three distinct locations within each stool specimen and transferred into 2 mL of RNAlater (Ambion, Austin, TX, USA) using collection tubes with an integrated sampling spoon (Sarstedt Sample Container, 15 ml; Nümbrecht, Germany). For metabolite analysis, fecal samples were collected from the central portion of the stool using the built-in sampling scoop of the collection container (Sarstedt Sample Container, 70 ml). Samples were transported to the laboratory at −20°C and subsequently stored at −80°C until further analysis.

### 2.3. 16S rRNA amplicon sequencing and sequence processing

16S rRNA gene amplicon sequencing was performed as previously described (Therdtatha et al., 2021). Briefly, DNA was extracted from stool samples using a bead-beating method and used to amplify the V3-V4 region of the 16S rRNA gene. Amplicons were indexed and sequenced on the Illumina MiSeq platform using the MiSeq Reagent Kit v3 (Illumina, San Diego, CA, USA).

Raw sequencing data were processed using the QIIME2 platform (Bolyen et al., 2019b;,Callahan et al., 2016) (version QIIME2-2024.5 for D-cohort and QIIME2-2021.2 for V-cohort). Sequence reads were quality-filtered and denoised and inferred as amplicon sequence variants (ASVs) using DADA2. Taxonomic assignment was performed using a scikit-learn naïve Bayes classifier (Bolyen et al., 2019b) trained on the SILVA database (release 138.2, SSU NR99) (Quast et al., 2012). The ASV tables for D-cohort (n =232 samples) and V-cohort (n = 404 samples) are provided in Supplementary Table S3 and S13, respectively.

### 2.4. SCFA analysis

In the D-cohort, approximately 100 mg of feces was suspended in 10 mM NaOH aqueous solution and centrifuged to obtain the supernatant. Crotonic acid was added to the supernatant to a final concentration of 10 mM as an internal standard. The supernatant was then mixed with chloroform, and the resulting aqueous phase was collected and subjected to organic acid analysis using a HPLC system (Shimadzu, Kyoto, Japan) as previously described (Dinoto et al., 2006).

In the V-cohort, approximately 100 mg of feces were thoroughly suspended in 700 μL of phosphate-buffered saline (PBS; 0.1 M, pH 7.4) prepared in deuterated water (MagniSov, Merk, Darmstadt, Germany) containing 4 mM sodium 3-(trimethylsilyl) propanoate-2,2,3,3,-*d*_4_ as an internal standard. After centrifugation to remove cell debris, the supernatant was subjected to quantitative NMR analysis as previously described (Therdtatha et al., 2021). SCFA concentrations for each sample are shown in Supplementary Tables S8 and S15 for D- and V-cohorts, respectively.

### 2.5. BA analysis

Sample preparation for BA analysis was performed according to the method described by Hagio et al (Hagio et al., 2009). Briefly, fecal samples were lyophilized, extracted with ethanol, partially purified, and subjected to LC-MS analysis. For the D-cohort, samples were analyzed using a Shimadzu LC-MS/MS 8050 system as described previously (Tanaka et al., 2020). Samples from the V-cohort were analyzed using the same instrument, with modified analytical conditions based on the Bile Acid Method Package Version 1 (Shimadzu). The quantified BA concentrations are provided in Supplementary Tables S8 and S15 for D- and V-cohorts, respectively.

### 2.6. Construction of co-abundance networks and identification of ASV-based CAGs

Detailed procedures for CAG construction, evaluation, and visualization are provided in Supplementary Note 2, including an overall workflow table (Supplementary Table S17). A total of 5,419,979 high-quality reads from the D-cohort were processed into 3,523 ASVs. After quality filtering and prevalence-based filtering, 1,189 ASVs were retained for downstream analyses.

Pairwise associations among retained ASVs were inferred using FastSpar (v1.0.0) (Watts et al., 2019), a compositionality-aware implementation of the SparCC algorithm (Friedman and Alm, 2012). Statistical significance and robustness of inferred correlations were evaluated using bootstrap resampling. Empirical p-values were adjusted for multiple testing using the Benjamini-Hochberg method (Benjamini and Hochberg, 1995).

A co-abundance network was constructed by retaining statistically significant associations after network sparsification to reduce spurious edges. The resulting weighted network was analyzed using the Louvain community detection algorithm (Blondel et al., 2008) as implemented in the igraph package (Csardi and Nepusz, n.d.), and the identified network modules were defined as co-abundant groups (CAGs). The resulting network was visualized using Cytoscape (version 3.10.3).

The robustness of CAG identification was evaluated under multiple clustering conditions using normalized mutual information (NMI), adjusted Rand index (ARI), and CAG module-size distributions (Supplementary Note 1). For downstream analyses, ASV abundances were aggregated within each CAG to generate a sample-by-CAG abundance matrix (Supplementary Table S6). Coverage metrics were calculated to evaluate how well the retained CAGs represented the original ASV dataset.

### 2.7. Association analyses of CAGs with SCFAs and BAs

Detailed procedures for the CAG-metabolite association analyses and visualization are provided in Supplementary Note 3. To evaluate the biological relevance of the identified CAGs, their associations with microbial metabolites were assessed using redundancy analysis (RDA). A sample-by-CAG matrix of relative abundances was used as the explanatory variable set, whereas concentrations of eight SCFAs and four metabolically defined BA groups were used as response variables. Both predictor and response matrices were centered and scaled to unit variance before analysis. RDA was performed using the vegan package in R, and statistical significance was assessed using permutation tests (999 permutations).

To further illustrate the hierarchical relationships among bacterial genera, CAGs, SCFAs, and BA metabolite groups, a Sankey diagram was constructed by integrating ASV taxonomic assignments with statistically significant CAG-metabolite associations identified by Spearman’s rank correlation analysis. Visualization was performed using the networkD3 package, with htmlwidgets and chromote used for SVG export. Links between genera and CAGs represent the taxonomic assignments of ASVs, whereas links between CAGs and metabolites represent statistically significant Spearman correlations (P < 0.05). Pairwise Spearman’s rank correlations between CAGs and metabolites were visualized as heatmaps.

### 2.8. CAG-based microbiome typing

The sample-by-CAG abundance matrix (Supplementary Table S6) was re-normalized to relative abundances, and pairwise Jensen-Shannon divergence (JSD) was calculated between samples. Partitioning around medoids (PAM) clustering was then applied across candidate cluster numbers ranging from K = 3 to K = 10 to identify gut microbiome types (GMTs). Candidate cluster numbers (K = 3-10) were evaluated using PAM clustering based on Jensen-Shannon divergence. The final GMT solution was selected by jointly considering average silhouette width, the explanatory power of each clustering solution for fecal short-chain fatty acid (SCFA) and bile acid (BA) profiles as assessed by the adjusted R² of redundancy analysis (RDA), and minimum cluster size. The robustness and validity of the selected GMT classification were evaluated using principal coordinate analysis (PCoA), permutational multivariate analysis of variance (PERMANOVA) (Anderson, 2017), betadisper analysis (Anderson et al., 2006), bootstrap resampling with stability assessed by the adjusted Rand index (ARI) and normalized mutual information (NMI), and cross-validation. Detailed procedures are described in Supplementary Note 4.

To facilitate interpretation of the identified GMTs, mean relative abundances of individual CAGs were calculated for each GMT, standardized by Z-score across GMTs, and visualized as a heatmap (Supplementary Figure S4).

To assess the temporal stability of GMTs, GMT assignments were compared between the two sampling time points (T0 and T1) for each subject. Subject-level transitions were visualized using a Sankey diagram generated with the ggalluvial package in R, and transition probabilities were calculated for each GMT at T0 as the proportion of subjects who remained in or transitioned to each GMT at T1. Detailed procedures are described in Supplementary Note 6.

### 2.9. Characterization of GMT-associated microbial and metabolic features

To characterize the biological features associated with each GMT, overall differences in fecal SCFA and BA profiles were first evaluated using permutational multivariate analysis of variance (PERMANOVA). Associations of GMTs with genus-level microbial composition, alpha diversity indices, individual SCFAs, and BAs were subsequently evaluated using MaAsLin2. (Mallick et al., 2021)

For MaAsLin2 analysis, a binary indicator variable representing membership in the target GMT (cluster_is_k) was generated, and one-versus-rest comparisons were performed across all samples. Linear models were fitted using cluster_is_k as a fixed effect (expr ∼ cluster_is_k), with no random effects included. No additional normalization or transformation was applied, and multiple testing correction was performed using the Benjamini-Hochberg false discovery rate (FDR) procedure. For genus-level analyses, only genera present in at least 10% of samples (≥24 of 232 samples) were retained for downstream analyses. Detailed procedures are described in Supplementary Note 5.

### 2.10. Associations of GMTs with host, dietary, and lifestyle factors in the D-cohort

Associations between gut microbiome types (GMTs) and host-related variables were evaluated using MaAsLin2 (Mallick et al., 2021). Analyses were restricted to individuals whose GMT assignments remained stable across the two sampling time points (n = 83). Variables were grouped into physiological characteristics, lifestyle factors, disease history, infection history, medication use, gastrointestinal characteristics, and dietary variables, including food groups, macronutrients, fatty acid intake, and micronutrients. Detailed descriptions of the variable groups are provided in Supplementary Note 7, and the variables included in each category are listed in Supplementary Tables S1 and S2. Each variable group was analyzed separately using the same analytical framework. For each GMT, one-versus-rest analyses were performed using GMT membership (target GMT versus all remaining GMTs) as the sole fixed effect (expr ∼ cluster_is_k). Continuous variables were standardized by z-score transformation, whereas no normalization or data transformation was applied. Associations were evaluated using linear models implemented in MaAsLin2, and P-values were adjusted for multiple testing using the Benjamini-Hochberg false discovery rate (FDR) procedure. Associations with q < 0.10 were considered statistically significant.

### 2.11. Application of the D-cohort CAG-GMT framework to V-cohort

Samples in the validation cohort were assigned to the predefined GMTs using the CAG-GMT framework established in the D-cohort. ASVs in the validation cohort were matched to ASVs assigned to the 27 CAGs identified in the D-cohort based on identical ASV sequences, and read counts were aggregated within each CAG (Supplementary Table S13). Aggregated counts were converted to relative abundances to generate sample-level relative abundance profiles of the 27 CAGs (Supplementary Table S16). Cluster centroids for each GMT were calculated by averaging the CLR-transformed CAG profiles of D-cohort samples assigned to each GMT after adding a pseudocount of 0.5 (Supplementary Table S18). The V-cohort CAG profiles were transformed using the same CLR procedure, and each sample was assigned to one of the seven predefined GMTs based on the nearest Euclidean distance to the corresponding centroid.

To evaluate whether the transferred GMT framework preserved CAG-based community structure across cohorts, PERMANOVA was performed on the combined D- and V-cohorts using Euclidean distances calculated from CLR-transformed CAG abundance profiles. GMT assignment and cohort were included as explanatory variables, and statistical significance was assessed using 999 permutations. The reproducibility of GMT-associated bile acid and SCFA profiles between the D- and V-cohorts was assessed using Mantel tests based on Bray-Curtis distance matrices.

The procedure and validation for applying the D-cohort CAG-GMT framework to the validation cohort is described in detail in Supplementary Note 8.

### 2.12. Association of GMTs with clinical parameters in V-cohort

Detailed procedures for statistical modeling and model evaluation are provided in Supplementary Notes 9. Associations between GMTs and host clinical parameters were evaluated using multinomial logistic regression, with GMT assignment as the outcome variable and a predefined clinical parameter panel as explanatory variables. The panel included body mass index (BMI), HbA1c, triglycerides (TG), high-density lipoprotein cholesterol (HDL), low-density lipoprotein cholesterol (LDL), γ-glutamyl transpeptidase (γ-GTP), alanine aminotransferase (ALT), estimated glomerular filtration rate (eGFR), uric acid (UA), systolic blood pressure (SBP), and diastolic blood pressure (DBP). Age, sex, and medication status were included as covariates. Model performance was evaluated using likelihood ratio tests, McFadden’s pseudo-R², Nagelkerke’s R², and five-fold cross-validation. In addition, distance-based redundancy analysis (dbRDA) was performed to assess the multivariate relationship between GMTs and host clinical biomarker profiles.

To further characterize GMT-specific associations with individual clinical parameters, binary logistic regression models were fitted for each clinical parameter using GMT assignment as the explanatory variable and age as a covariate. Adjusted probabilities for abnormal clinical parameters were estimated as predictive margins with 95% confidence intervals (CIs) using the postestimation margins command in Stata/SE 12 (StataCorp, College Station, TX, USA). Abnormal values were defined according to standard reference ranges used in routine health check-ups in Japan (Supplementary Table S12).

### 2.13. Comparison of GMTs and representative genera for clinical associations

To further evaluate the transferability and generalizability of the CAG-based GMT framework, its performance was compared with that of a de novo genus-based microbiome stratification generated independently from the V-cohort. Genus-based microbiome types were generated by de novo clustering of genus-level relative abundance profiles from the V-cohort using the same clustering procedure applied to the D-cohort. The resulting genus-based microbiome types were evaluated using the same multinomial logistic regression framework applied to the transferred CAG-based GMTs, and model performance was compared using likelihood ratio tests, McFadden’s pseudo-R², Nagelkerke’s R², and five-fold cross-validation.

To compare the explanatory value of GMTs with that of representative genera, additional Firth logistic regression analyses were performed for selected GMT-clinical associations. MT-4 and MT-5 were selected because they showed the strongest and most reproducible associations with liver and renal function, respectively. *Faecalibacterium* and *Bifidobacterium*, which consistently characterized MT-4 and MT-5, respectively, in both the D- and V-cohorts, were selected as representative genera. For each GMT-clinical association, three Firth logistic regression models were fitted after adjustment for age, sex, and medication use: (i) an MT-only model, (ii) a genus-only model, and (iii) a combined model including both the GMT and its corresponding representative genus. The additional explanatory value of GMTs and representative genera was evaluated using nested likelihood ratio tests comparing the corresponding regression models. Detailed procedures are described in Supplementary Note 10.

### 2.12. Ethics approval and consent to participate

The protocol for D-cohort was approved by the ethics committees of the Faculty of Agriculture, Kyushu University (No. 25, 41, 48) and Ajinomoto Co., Inc. (No. 2015-030). The protocol for V-cohort was approved by the ethics committees of Tohoku Chuo Hospital (Nos. 201-3 and 210-6), Kyushu University, Faculty of Agriculture (No. 78), and Nagoya City University (No. 834-6). Written informed consent was obtained from all participants prior to enrollment.

## 3. RESULTS

### 3.1. Inter-individual and temporal variability of the gut microbiome in the D-cohort

A total of 3,523 ASVs were inferred from 5,419,979 high-quality 16S rRNA V3-V4 sequences obtained from 232 fecal samples of 116 subjects, and taxonomic profiles were assigned from the phylum to species level (Supplementary Table S3). Consistent with previous Japanese cohort studies (Nishijima et al., 2016; Odamaki et al., 2016; Takagi et al., 2022; Watanabe et al., 2021), a broad trade-off relationship was observed between the two dominant phyla, Bacillota (formerly known as Firmicutes) and Bacteroidota (formerly known as Bacteroidetes), while temporal changes over the 1-month interval were generally modest (Figure 2A, left). Moreover, the subdominant phylum Actinomycetota (formerly known as Actinobacteria) showed substantial inter-individual variation with its major genus *Bifidobacterium* ranging from 0 to 45.3%.

**Figure 2.**
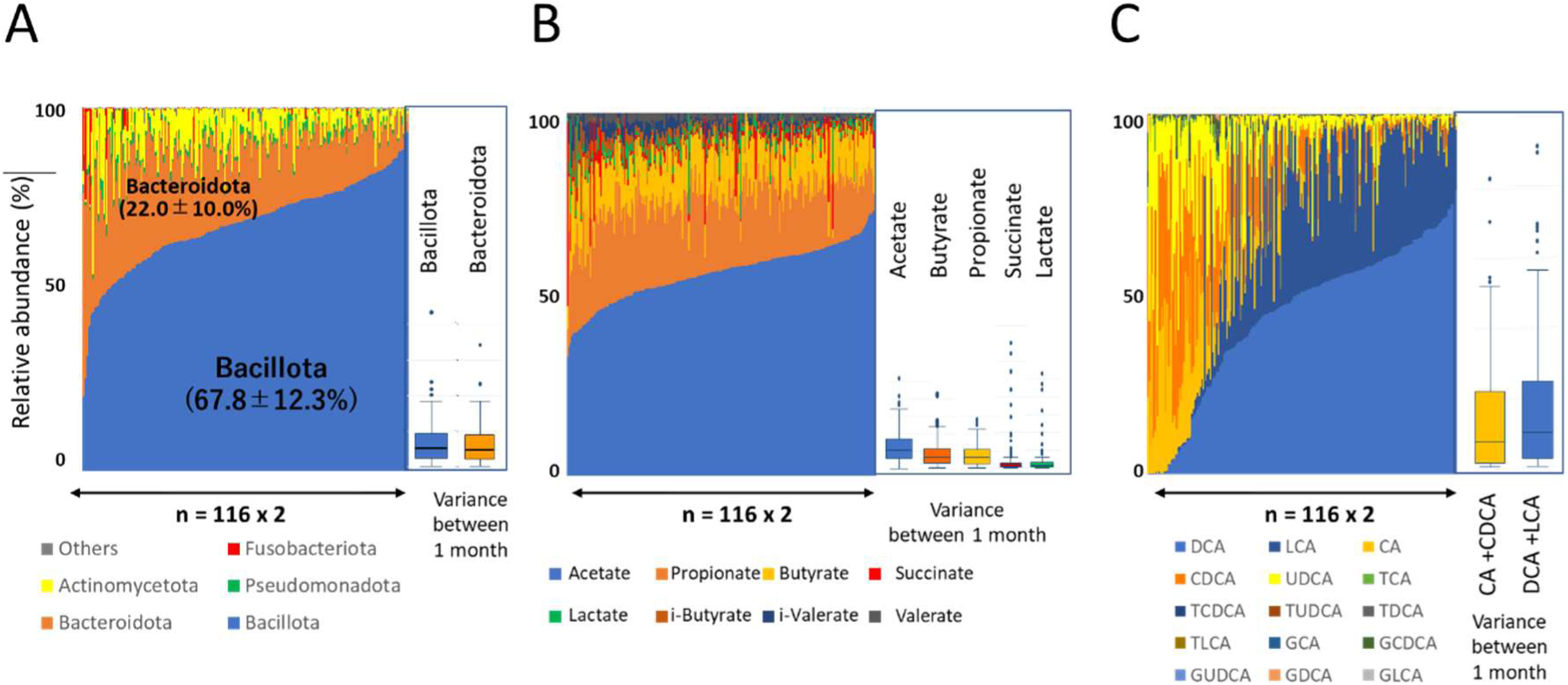
Inter-individual and temporal variation in phylum-level fecal microbiome (A), SCFA (B), and BA (C) profiles. In each panel, the relative abundance profile of each sample is shown as a stacked bar plot. The 232 samples collected from 116 individuals at a one-month interval are ordered according to the abundance of the dominant component in each panel (Bacillota in A, acetate in B, and DCA in C). Box plots adjacent to each stacked bar plot summarize temporal variation in the dominant components between the two sampling time points.

SCFA profiles were dominated by acetate, followed by propionate and butyrate, and were likewise relatively stable over the 1-month interval (Figure 2B). In contrast, BA composition showed substantially greater inter-individual and temporal variability, with the balance between primary and secondary BAs spanning nearly the full range across subjects (Figure 2C), indicating that BA metabolism appears to be a more dynamic aspect of the gut microbial ecosystem than community structure or SCFA metabolism.

Together, these observations highlight the structural and functional complexity of the gut microbial ecosystem, motivating subsequent CAG-based analyses to resolve this complexity through the delineation of ecological guilds.

### 3.2. Identification of CAGs in the D-cohort

Pairwise co-abundance relationships were inferred from 1,189 filtered ASVs (prevalence ≥1% and abundance ≥0.05%) using FastSpar. The resulting sparsified backbone network comprised 316 ASVs connected by 707 co-abundance edges exhibiting r > 0.2 (Figure 3A). Community detection analysis of this network identified 27 CAGs (Figure 3B). The identified CAGs spanned 32 families and 91 genera (Supplementary Table S5). Although the 316 CAG-associated ASVs represented only 8.97% of the total 3,524 detected ASVs, they accounted for 75.9% of total abundance and captured 79.5% of the total variance in ASV abundances across samples.

**Figure 3.**
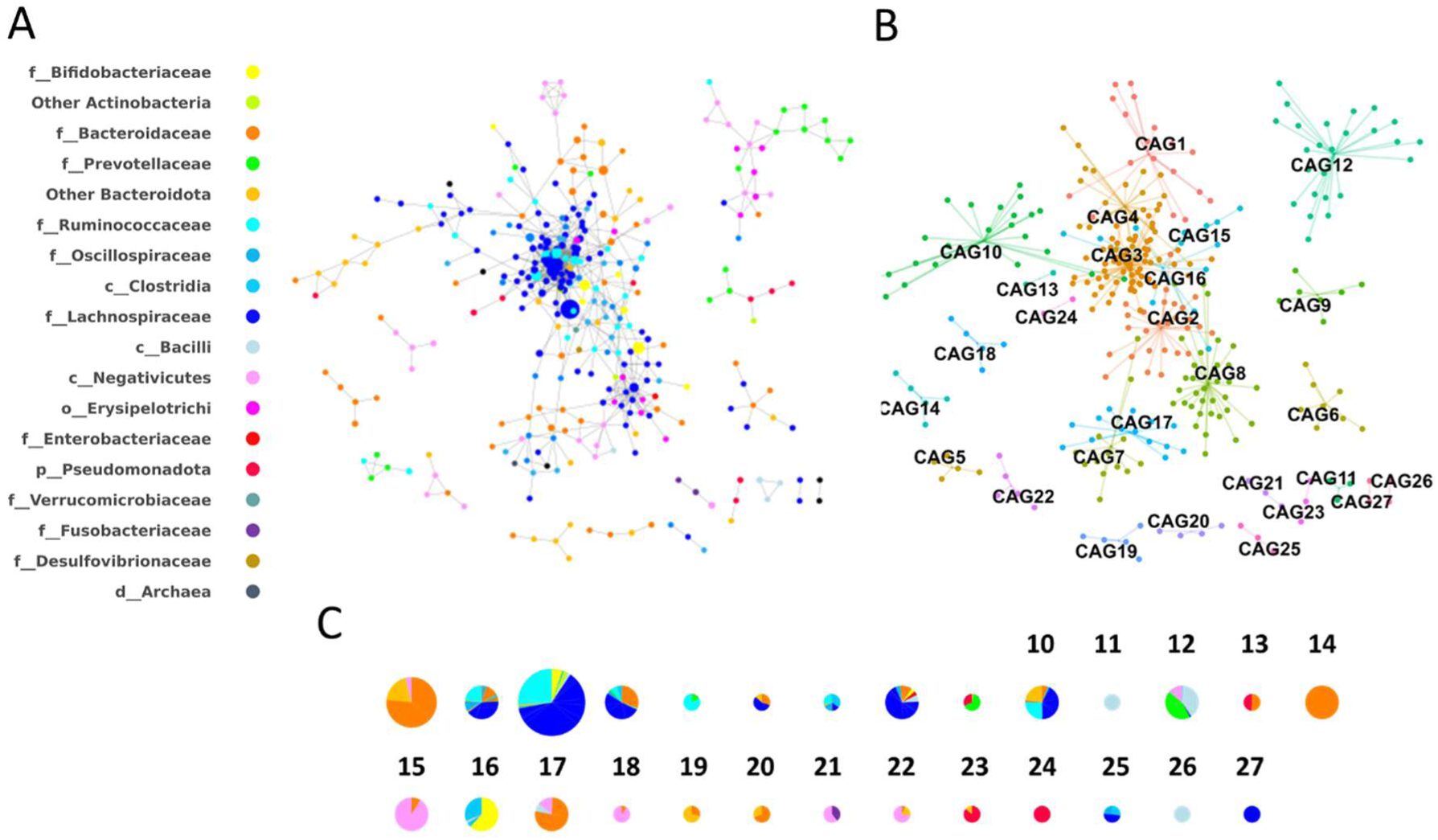
CAG-based microbiome typing of 232 samples in the discovery cohort. (A) Co-abundance network of 432 amplicon sequence variants (ASVs) shared by the 232 samples. The colors of the nodes represent taxonomic groups (family level or higher), as indicated in the taxonomic legend on the left. Node size is proportional to ASV abundance. (B) Co-abundance-defined ASV guilds. ASVs were clustered into co-abundance groups (CAGs) based on pairwise associations, and ASV nodes belonging to the same CAG are connected by edges of a unified color. (C) Pie charts showing the bacterial family composition of each CAG. Colors represent bacterial families and are the same as those in (A). Pie diameter represents the mean relative abundance of each CAG across the D-cohort.

To assess robustness, Louvain community detection was repeated using five different random seeds. The set of included ASVs remained identical across runs, with variability confined to minor differences in module boundaries. Pairwise comparisons across runs yielded a mean normalized mutual information (NMI) of 0.90 and a mean adjusted Rand index (ARI) of 0.78, demonstrating high reproducibility of the inferred CAG structure (Supplementary Table S19). The number of detected CAG modules varied only slightly, ranging from 26 to 27 across runs. In addition, highly similar CAG module-size distributions were observed across clustering solutions (Supplementary Table S20).

### 3.3. Structural organization and metabolic associations of CAGs

CAG size ranged from 2 to 61 ASVs, indicating substantial heterogeneity in module size and structure across the network (Figure 3B, Supplementary Table S4). Smaller CAGs generally exhibited higher network density, whereas larger CAGs tended to be more sparsely connected. Most CAGs comprised complex taxonomic assemblages rather than taxonomically homogeneous modules (Figure 3C). Within this taxonomic complexity, two major compositional patterns were apparent. Several large CAGs (e.g., CAG2, CAG3, CAG8) were dominated by members of Bacillota, whereas another group of CAGs (e.g., CAG1, CAG14, and CAG17) was characterized predominantly by members of Bacteroidota. In contrast, several smaller CAGs were composed almost exclusively of a single taxonomic group, including *Streptococcus* (CAG11) and *Sutterella* (CAG24). Other CAGs consisted of distinct combinations of phylogenetically distant taxa, including Bacillota-*Parabacteroides* (CAG10), *Segatella*-Erysipelatoclostridiaceae (CAG12) and *Bifidobacterium-*Peptostreptococcaceae (CAG16). These observations indicate that CAG organization reflects recurrent ecological co-abundance relationships rather than taxonomic relatedness alone.

To visualize the hierarchical organization linking ASVs, CAGs, SCFAs, and BAs, a Sankey diagram was constructed (Figure 4A). Two major ecological-metabolic architectures were apparent. One architecture was centered on the largest module, CAG3, which comprised diverse members of Bacillota and linked predominantly to butyrate production and secondary BA metabolism. CAG3 was highly interconnected and accounted for nearly half of all network edges (Figure 3B and Supplementary Table S4), suggesting that it represents a functional core microbiome module in the gut microbial community of this Japanese population. In contrast, ASVs belonging primarily to Bacteroidota were distributed across several CAGs that converged predominantly on propionate metabolism (Figure 4). Several of these CAGs were additionally connected with valerate. Within this Bacteroidota-associated architecture, comparatively fewer ASV connections extended toward BA metabolite groups. These global CAG-metabolite associations were quantitatively evaluated by RDA, which demonstrated significant associations of CAG profiles with both SCFA and BA profiles, whereas genus-level profiles showed weaker statistical support for SCFA profiles and no significant association with BA groups (Table 2). These findings indicate that CAG-based organization captures microbial metabolic associations more consistently than genus-level classification.

**Figure 4.**
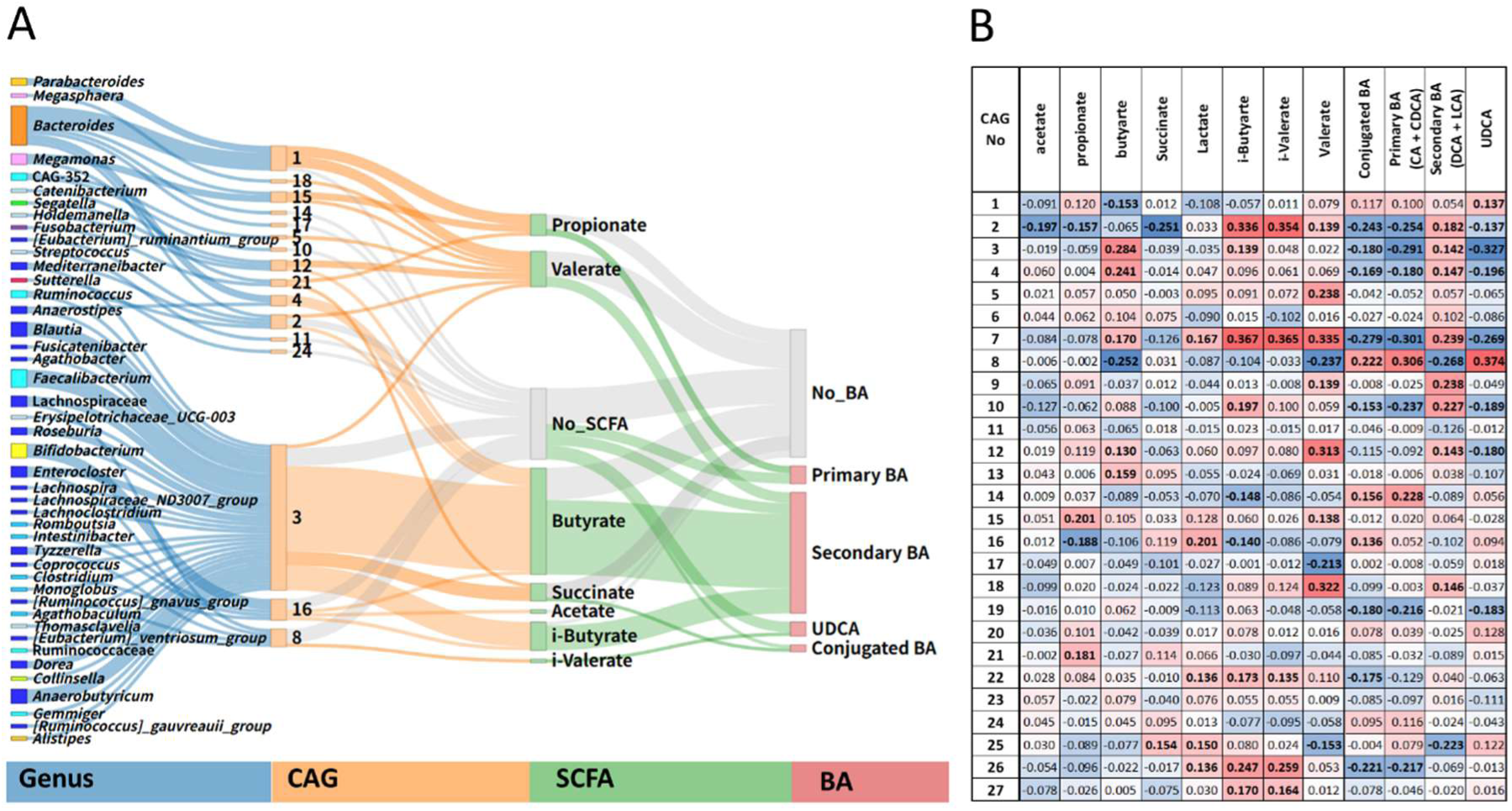
Association profiles of CAGs with SCFAs and BAs in the D-cohort. (A) Sankey diagram illustrating the hierarchical organization linking bacterial genera, CAGs, fecal short-chain fatty acids (SCFAs), and bile acid (BA) metabolite groups. Each ASV was assigned to both a genus and a CAG, and CAG-metabolite connections represent statistically significant associations identified by Spearman correlation analysis. Ribbon width is proportional to the number of ASVs represented by each connection. Blue, orange, and green ribbons denote connections originating from the genus, CAG, and SCFA layers, respectively. Colors of the genus boxes indicate taxonomic groups (family level or higher), as defined in Figure 3A. Gray ribbons leading to **No_SCFA** or **No_BA** indicate ASVs belonging to CAGs that showed no significant association with any measured SCFA or BA metabolite group, respectively. BA metabolite groups were defined as follows: conjugated BA (GCA, TCA, GCDCA, TCDCA, GDCA, TDCA, GLCA, TLCA, GUDCA, and TUDCA), primary BA (CA and CDCA), and secondary BA (DCA and LCA). UDCA is shown separately. **(B)** Heatmap showing Spearman correlation coefficients between individual CAGs and fecal metabolites. Positive and negative correlations are indicated by red and blue shading, respectively. Bold numbers indicate significant correlations (*P* < 0.05). Metabolites are arranged in the same order as in panel A to facilitate comparison between the global hierarchical architecture and individual CAG-metabolite associations.

**Table 2.** Comparison of genus- and CAG-based RDA models for fecal SCFA and BA profiles.

| Microbiome feature | Metabolite profile * | Adjusted R<sup>2</sup> † | Global P ‡ |
| --- | --- | --- | --- |
| Genera | SCFAs | 0.281 | 0.011 |
| Genera | BA groups | 0.143 | 0.233 |
| CAGs | SCFA | 0.160 | 0.001 |
| CAGs | BA groups | 0.163 | 0.001 |

\*The same SCFA and BA variables as those listed in Table 3 were used for each RDA model.
†Adjusted R<sup>2</sup> indicates the explanatory power of each RDA model.
‡Global P values were obtained by permutation testing (999 permutations).

Individual CAG-metabolite associations were then examined using Spearman correlation analysis (Figure 4B). SCFA associations exhibited distinct metabolite-specific patterns. Several Bacillota-enriched CAGs, including CAG3, CAG4, and CAG7, showed positive associations with butyrate, whereas propionate was associated primarily with CAG15, a module enriched in *Bacteroides* and *Megamonas*. Acetate showed no preferential association with any individual CAG. The *Prevotellaceae*-enriched CAG12 was specifically associated with valerate. Further, CAG2 and CAG7, enriched in Christensenellaceae, showed marked association with branched-chain fatty acids (BCFAs).

Associations between CAGs and BA metabolite groups formed two broad patterns: a secondary (7α-dehydroxylated) BA-associated type and a conjugated/primary BA-UDCA-associated type (Figure 4B). CAG2, CAG3, CAG4, CAG7, CAG10, and CAG12 were associated predominantly with secondary BAs, whereas CAG1, CAG8, and CAG14 were associated primarily with conjugated and primary BAs together with UDCA. Notably, CAG8 exhibited the strongest positive association with UDCA.

### 3.4. CAG-based microbiome typing in the D-cohort

To stratify gut microbial community structure in the D-cohort, we performed unsupervised clustering of 232 samples based on the relative abundance profiles of 27 co-abundance groups (CAGs). Two clustering approaches were compared: partitioning around medoids based on Jensen-Shannon divergence (PAM-JSD) and hierarchical clustering using Ward’s method with Aitchison distance (WARD-AIT).

The optimal number of gut microbiome types (GMTs) was determined by integrating multiple criteria. Average silhouette widths did not identify a distinct optimum across the tested range of K values. In contrast, the explanatory power of the clustering solutions for fecal SCFA and BA profiles, evaluated by the adjusted R² of redundancy analysis (RDA), increased with K before reaching a plateau (Supplementary Figure S1). Considering the trade-off between explanatory power and cluster size (Supplementary Table S21), while requiring each cluster to contain at least 10 samples, K = 7 was selected as the optimal solution for the PAM-JSD method. Accordingly, the D-cohort was classified into seven gut microbiome types (MT-1 to MT-7).

Although the GMTs were not completely separated in two-dimensional PCoA space (Supplementary Figure S2), PERMANOVA demonstrated significant differences among clusters (R² = 0.501, P = 0.001), indicating that the clustering captured substantial variation in microbiome structure. Although betadisper also detected significant differences in within-cluster dispersion (P = 2.48 × 10⁻⁵), the separation of cluster centroids in ordination space together with the PERMANOVA results suggests that the observed clustering reflects differences in community composition rather than dispersion alone. Cross-validation and bootstrap analyses further supported the stability of the clustering (mean cross-validation accuracy = 0.67, mean bootstrap ARI = 0.47, and mean bootstrap NMI = 0.59; Supplementary Figure S3). Together, these results indicate that the GMT framework captures robust ecological structure within the gut microbiome while preserving gradual transitions between community states rather than defining completely discrete groups.

The identified GMTs exhibited distinct CAG enrichment patterns (Supplementary Figure S4). Several CAGs showed preferential enrichment in specific GMTs, indicating that each microbiome type was characterized by a unique configuration of co-abundance structures rather than by enrichment of a single dominant module. These CAG configurations were accompanied by distinct genus-level compositions (Figure 5A), providing a taxonomic interpretation of the underlying co-abundance patterns.

**Figure 5.**
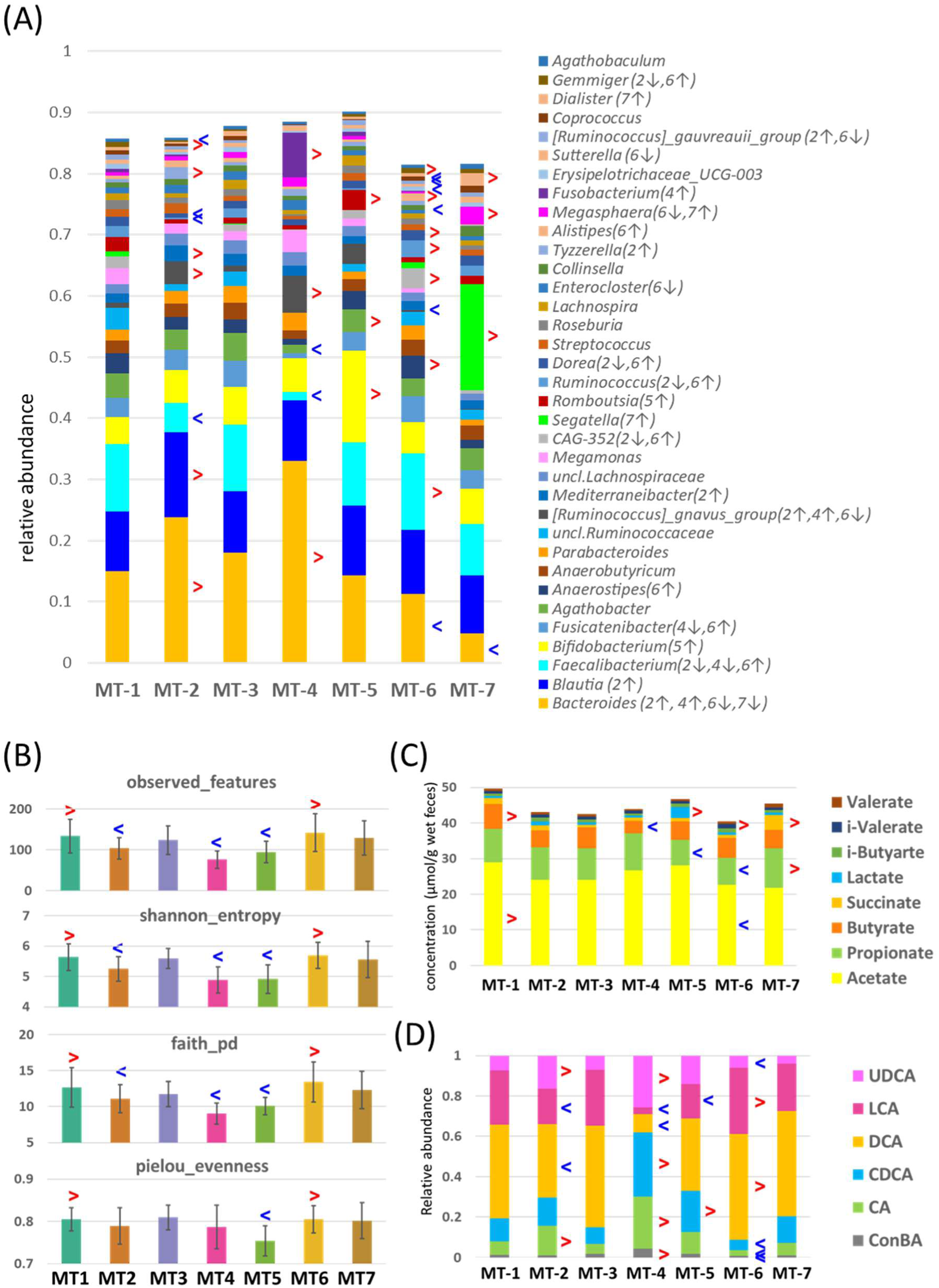
Characteristics of the seven GMTs in the D-cohort. (A) Genus-level composition. Stacked bar plots showing the mean relative abundances of bacterial genera with an overall mean relative abundance ≥0.5% in the discovery cohort (D-cohort). Genera are ordered from bottom to top according to their overall mean relative abundances. Numbers in parentheses following each genus name indicate the GMT(s) in which the genus was significantly enriched (↑) or depleted (↓). Differential abundance analysis was restricted to genera detected in at least 10% of samples (≥24 of 232 samples). (B) Mean alpha diversity indices (Observed features, Shannon entropy, Faith’s phylogenetic diversity, and Pielou’s evenness) for each GMT. Error bars indicate standard deviations (SD). (C) Stacked bar plots showing the mean fecal concentrations of major short-chain fatty acids (SCFAs) across the seven GMTs. (D) Stacked bar plots showing the mean relative composition of major fecal bile acids (BAs) across the seven GMTs. In all panels, red and blue arrowheads indicate features significantly enriched or depleted, respectively, in each GMT compared with the remaining GMTs, as determined by one-versus-rest analysis using MaAsLin2 with Benjamini-Hochberg false discovery rate correction (q ≤ 0.10).

### 3.5. Characterization of GMT structure and taxonomy

The seven GMTs exhibited distinct taxonomic compositions and ecological characteristics (Figure 5). Taxonomically (Figure 5A), MT-4 was distinguished by marked enrichment of *Bacteroides*, the *Ruminococcus gnavus* group, and *Fusobacterium*, together with depletion of *Faecalibacterium*. MT-2 shared several taxonomic characteristics with MT-4, although these features were less pronounced. MT-5 was characterized by enrichment of *Bifidobacterium* and *Romboutsia*, whereas MT-6 exhibited depletion of *Bifidobacterium* but enrichment of *Faecalibacterium* and *Anaerostipes*. MT-7 was characterized by enrichment of *Segatella* and *Megasphaera* together with depletion of *Bacteroides*. In contrast, MT-1 and MT-3 displayed more balanced taxonomic compositions, characterized by relatively even abundances of the three dominant genera, *Bacteroides*, *Blautia*, and *Faecalibacterium*. Alpha diversity was lower in MT-4 and MT-5 than in the other GMTs, while MT-1 and MT-6 showed relatively high alpha diversity (Figure 5B). Notably, the lower alpha diversity in MT-4 coincided with marked taxonomic shifts.

### 3.6. Association of GMTs with SCFA and BA profiles

To evaluate the functional relevance of the identified GMTs, we examined their associations with fecal SCFA and BA profiles using PERMANOVA (Table 3). GMT classification was significantly associated with both metabolite classes. SCFA profiles differed significantly among GMTs, although with a modest effect size (R² = 0.110, F = 4.637, P = 0.001). Likewise, individual bile acid (BA) profiles showed significant but modest separation across GMTs (R² = 0.117, F = 4.960, P = 0.001). To better capture biologically meaningful variation in bile acid metabolism, BAs were further grouped into four functional categories comprising primary BAs (CA and CDCA), secondary BAs (DCA and LCA), UDCA, and conjugated BAs. PERMANOVA based on these categorized BA profiles yielded a substantially larger effect size (R² = 0.234, F = 11.43, P = 0.001) than analyses based on individual BA species, indicating that biologically grouped BA profiles more effectively captured functional differences among GMTs.

**Table 3.** PERMANOVA showing differences in fecal SCFA and BA profiles among GMTs.

| Parameter Category | Variables (n) | R<sup>2</sup> | F | p-value |
| --- | --- | --- | --- | --- |
| SCFA<sup>a</sup> | 8 | 0.110 | 4.637 | 0.001 |
| Bile acids (individual)<sup>b*</sup> | 15 | 0.117 | 4.960 | 0.001 |
| Bile acids (categorical)<sup>c*</sup> | 4 | 0.234 | 11.431 | 0.001 |

<sup>a</sup>SCFA profile comprised acetate, propionate, butyrate, succinate, lactate, isobutyrate, isovalerate, and valerate.
<sup>b</sup>Individual BA profile comprised CA, CDCA, UDCA, DCA, LCA, TCA, TCDCA, TUDCA, TDCA, TLCA, GCA, GCDCA, GUDCA, GDCA, and GLCA.
<sup>c</sup>BA category profile comprised primary bile acids (CA + CDCA), secondary bile acids (DCA + LCA), UDCA, and conjugated bile acids.
\*Relative abundances (proportions of total bile acids) were used for all bile acid analyses.

The metabolite profiles associated with each GMT are summarized in Figure 5C, Figure 5D, and Supplementary Table S10. MT-4 exhibited elevated levels of primary BAs (CA and CDCA) together with increased conjugated BAs, whereas MT-6 showed comparatively lower levels of both primary and conjugated BAs. These contrasting BA profiles were consistent with the distribution of CAG3, the largest core CAG identified in the co-abundance network and a major CAG positively associated with secondary BAs (Fig. 4A), suggesting its potential involvement in the 7α-dehydroxylation of primary BAs.

SCFA profiles likewise differed among GMTs. Among the three major SCFAs, MT-6 exhibited significantly higher butyrate levels, whereas MT-7 showed significantly higher propionate levels. In contrast, MT-4 exhibited significantly lower butyrate levels. Intermediate metabolites also displayed GMT-specific patterns, with MT-5 showing elevated lactate levels and MT-7 exhibiting increased succinate levels. Branched-chain fatty acid metabolism differed among GMTs as well, with MT-6 showing increased i-butyrate and MT-7 exhibiting higher valerate levels. These metabolite signatures were broadly consistent with the microbial characteristics of the corresponding GMTs. Elevated butyrate in MT-6 coincided with enrichment of *Faecalibacterium* and *Anaerostipes*, two well-recognized butyrate producers, whereas increased propionate and succinate in MT-7 were consistent with enrichment of *Segatella*, a genus associated with succinate-dependent propionate production. Likewise, elevated lactate in MT-5 corresponded to enrichment of *Bifidobacterium*.

### 3.7. Stability of GMTs over time

To evaluate the temporal stability of GMTs, GMT assignments at the two sampling time points were compared using a Sankey plot (Figure 6, Supplementary Table S7). Overall, 83 of the 116 subjects retained the same GMT across the two time points, although stability varied among GMTs, ranging from 52.6% for MT3 to 88.2% for MT2. Notably, GMT transitions were not random but were largely confined to a limited number of related GMTs, whereas transitions between more distinct GMTs were rarely observed.

**Figure 6.**
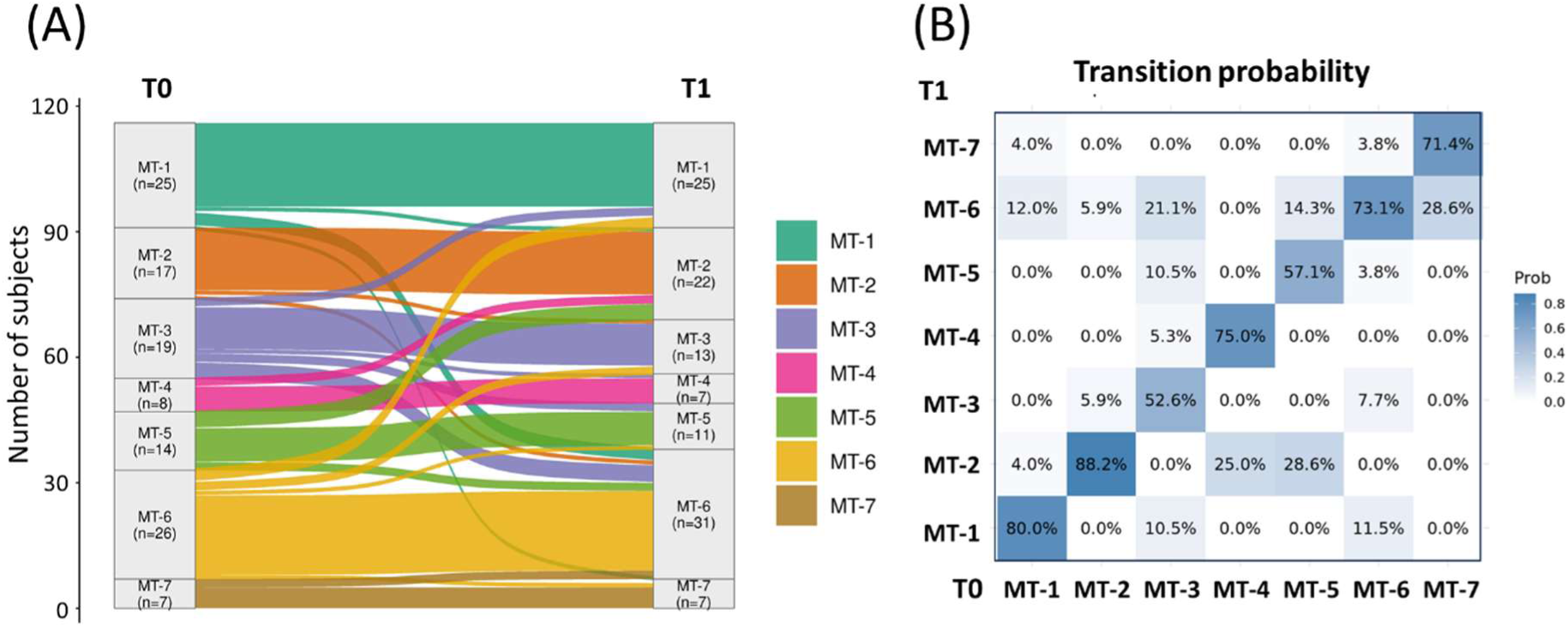
Temporal stability and transitions of GMTs over a 1-month interval. **(A)** Sankey diagram illustrating subject-level transitions between GMTs from T0 to T1. Flow widths are proportional to the number of subjects undergoing each transition. **(B)** Heatmap showing row-wise transition probabilities between GMTs. Each value represents the proportion of subjects assigned to a given GMT at T0 who were assigned to each GMT at T1.

### 3.8. Associations of GMTs with host and dietary factors

To comprehensively characterize host factors associated with GMTs, multivariable MaAsLin2 analyses were performed using subjects whose GMT assignments remained stable over the 1-month interval (n = 83). A broad range of host variables was evaluated, including anthropometric measurements, lifestyle factors, medical history, medication use, gastrointestinal symptoms, dietary intake assessed using a comprehensive dietary history questionnaire, and nutrient intake estimated from these dietary records (Supplementary Tables S1-S2). Detailed analytical procedures are described in the Materials and Methods and Supplementary Note 7. Overall, relatively few host-related variables showed significant associations with individual GMTs after FDR correction (q < 0.10). Significant associations were observed for age (MT-2), sex (MT-2), BMI (MT-4), physical activity (MT-1), probiotic use (MT-6), and alcohol consumption (MT-4) (Supplementary Table S22). Among these, the association between MT-4 and alcohol consumption was the most biologically plausible, whereas the overall host-associated differences among GMTs were modest.

### 3.9. Application of the CAG-based framework to V-cohort

To assess the reproducibility of the CAG-based framework, we analyzed an independent validation cohort (V-cohort) of 404 Japanese adults. A total of 12,025,774 sequencing reads corresponding to 8,143 ASVs were obtained. Of these, 304 ASVs, representing 8,797,084 reads, were assigned to the 27 CAGs defined in the D-cohort, accounting for 73.0% ± 10.4% of total reads per sample (Supplementary Table S13). The overall distribution of the 27 CAGs in the V-cohort was highly concordant with that in the D-cohort (Pearson’s r = 0.93, P < 0.001), indicating that the CAG framework identified in the D-cohort was broadly preserved in the V-cohort (Supplementary Fig. S5).

To further evaluate whether the functional characteristics of the CAGs were preserved across cohorts, association coefficients between individual CAGs and microbial metabolites were compared between the D- and V-cohorts. Significant positive correlations were observed for both SCFAs (R = 0.313, P < 0.001, adjusted R² = 0.137) and BAs (R = 0.754, P < 0.001, adjusted R² = 0.574), demonstrating that the metabolic characteristics associated with individual CAGs were largely conserved between the two cohorts, particularly for BA metabolism (Supplementary Figure S6).

The CAG-metabolite associations and their comparison with D-cohort are shown in Supplementary Figure S7. Overall, SCFA-associated CAG patterns were well preserved, although acetate, lactate, and succinate showed weaker reproducibility between cohorts, likely reflecting greater susceptibility to extrinsic factors than to CAG-associated microbial functions. Representative examples included the positive association of the Bacillota-dominant CAG3 with butyrate, whereas the *Prevotella*-dominant CAG12 and *Bacteroides-*dominant CAG15 were associated with higher propionate levels. Two major BA association patterns were observed in the validation cohort, mirroring those identified in the D-cohort. Specifically, CAG2, CAG3, CAG4, CAG7, CAG10, and CAG12 were associated predominantly with secondary BAs, whereas CAG1, CAG8, and CAG14 were associated primarily with conjugated and primary BAs together with UDCA. In addition, the V-cohort included quantitative measurements of LCA-derived metabolites, including isoalloLCA, 3-oxoLCA, and isoLCA, which also showed positive associations with the secondary BA-associated CAGs.

### 3.10. V-cohort microbiome typing using the 27-CAG framework

Based on the 27-CAG abundance profiles generated using the D-cohort framework (Supplementary Table S18), samples from the independent validation cohort were assigned to the seven predefined GMTs (MT1-MT7) using a centroid-based approach in CLR-transformed space. PERMANOVA analysis demonstrated that GMT identity explained a substantial proportion of the variance in microbial community composition (R² = 0.454, P = 0.001), indicating that the transferred GMT configuration was well preserved in the validation cohort (Supplementary Table S23).

Assignment confidence was generally high, with no samples showing negative assignment margins and a median margin of 0.071 (Supplementary Table S24). Silhouette analysis yielded a modest mean silhouette coefficient (∼0.14), consistent with the continuous nature of gut microbiome variation (Supplementary Table S25).

The overall CAG abundance patterns defining each GMT were highly consistent between the D- and V-cohorts, supporting the transferability of the 27-CAG-based microbiome typing framework to an independent population (Supplementary Figure S8). Centroid-based comparison between cohorts revealed moderate to strong concordance of GMT-specific CAG composition profiles (Pearson’s r = 0.628-0.809), with the dysbiosis-associated MT-4 showing the highest concordance (r = 0.809) (Supplementary Table S26). Consistent with these observations, PERMANOVA including both GMT and cohort effects confirmed that GMT assignment accounted for the majority of variation in CAG composition (R² = 0.451), whereas cohort effects were negligible (R² = 0.0037) (Supplementary Table S27).

Taxonomic characteristics of each GMT in the validation cohort were broadly consistent with those observed in the D-cohort (Figure 7A). In particular, the dysbiosis-like signature of MT-4, characterized by depletion of *Faecalibacterium*, enrichment of *Fusobacterium*, and *Ruminococcus gnavus* group and reduced alpha diversity (Figure 7B), was well preserved in the validation cohort. Characteristic taxonomic features of MT-5 and MT-7, including enrichment of *Bifidobacterium* and *Segatella*, respectively, were also maintained. These observations indicate that the major taxonomic characteristics defining individual GMTs were reproducible across independent cohorts.

**Figure 7.**
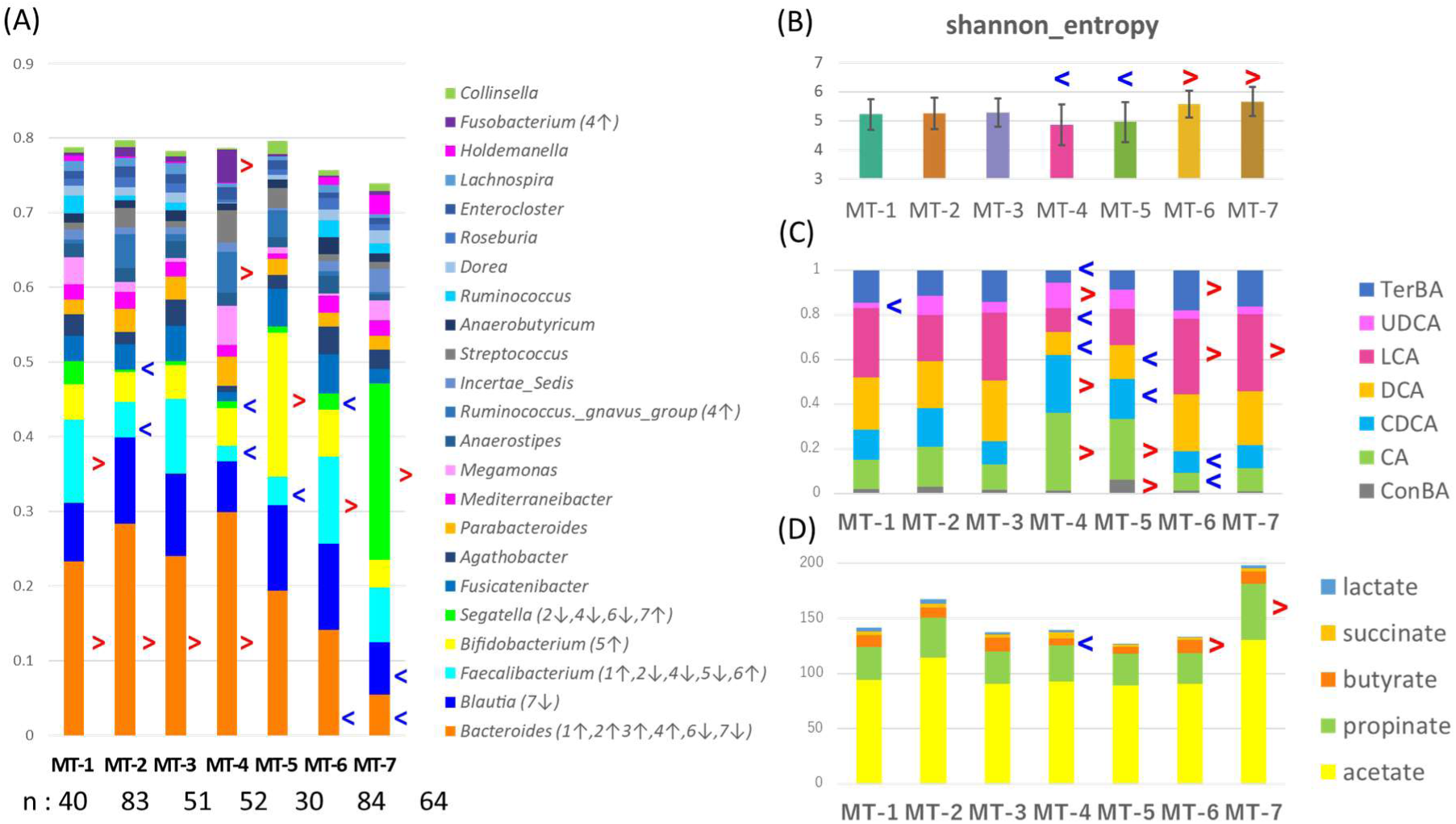
Characteristics of the seven GMTs in the V-cohort. (A) Genus-level composition. Stacked bar plots showing the mean relative abundances of bacterial genera with an overall mean relative abundance ≥5% in the validation cohort. Genera are ordered from bottom to top according to their overall mean relative abundances. Numbers in parentheses following each genus name indicate the GMT(s) in which the genus was significantly enriched (↑) or depleted (↓). For clarity, only the ten most significantly enriched and depleted genera are indicated. (B) Mean Shannon entropy for each GMT. Error bars indicate standard deviations (SD). (C) Stacked bar plots showing the mean fecal concentrations of major short-chain fatty acids (SCFAs) across the seven GMTs. (D) Stacked bar plots showing the mean relative composition of major fecal bile acids (BAs) across the seven GMTs. In all panels, red and blue arrowheads indicate features significantly enriched or depleted, respectively, in each GMT compared with the remaining GMTs, as determined by one-versus-rest analysis using MaAsLin2 with Benjamini-Hochberg false discovery rate correction (q ≤ 0.10).

Functional metabolite profiles also showed broadly consistent patterns across cohorts. Lower secondary bile acid ratios in MT-4 and MT-5 and higher secondary bile acid ratios in MT-6 and MT-7 were reproducibly observed, although not all differences reached statistical significance (Figure 7C). Consistent with these observations, the overall GMT-level BA composition was strongly preserved between cohorts, as demonstrated by Mantel analysis (Mantel r = 0.869, P < 0.001). SCFA profiles likewise showed similar trends, particularly reduced butyrate in MT-4 and elevated propionate in MT-7 (Figure 7D). However, preservation of the overall GMT-level SCFA structure did not reach statistical significance (Mantel r = 0.313, P = 0.168), indicating greater inter-cohort variability in SCFA profiles.

Taken together, these findings demonstrate that the GMTs identified in the D-cohort were reproducibly represented in the V-cohort, preserving both taxonomic and metabolic characteristics across independent cohorts.

### 3.11. Associations between GMTs and clinical and lifestyle characteristics

To evaluate the relationship between GMTs and host clinical status, we first examined the multivariate structure of the clinical biomarker panel using distance-based redundancy analysis. Clinical biomarker profiles showed a structured distribution across samples, with the constrained ordination model explaining 17.7% of the adjusted variation (Supplementary Figure S9). Although GMTs were not clearly separated in the ordination space, GMT was significantly associated with variation in host clinical profiles (permutation test, P = 0.007).

To determine whether these associations were independent of demographic factors, we performed multinomial logistic regression using a predefined clinical panel while adjusting for age, sex, and medication use (Supplementary Table S28). Adding the clinical panel significantly improved model fit compared with the demographic-adjusted model (McFadden’s pseudo-R² = 0.092 vs. 0.023; Nagelkerke’s R² = 0.298), with a significant incremental contribution of the clinical variables (ΔMcFadden’s pseudo-R² = 0.070; likelihood ratio test, P = 0.002). Consistently, the clinical panel remained significantly associated with GMT assignment after adjustment for medication alone (likelihood ratio test, P < 0.001; McFadden’s pseudo-R² = 0.081; Nagelkerke’s R² = 0.266), indicating that clinical parameters explained GMT variation beyond demographic factors and medication use.

We then examined GMT-specific patterns in clinical parameters using adjusted probabilities derived from logistic regression models (Fig. 8A, B). Significant GMT-specific associations were observed for several clinical parameters. Multinomial logistic regression further showed a significant overall association between the clinical panel and GMT assignment (likelihood ratio test, P = 0.002; Supplementary Table S28).

**Figure 8.**
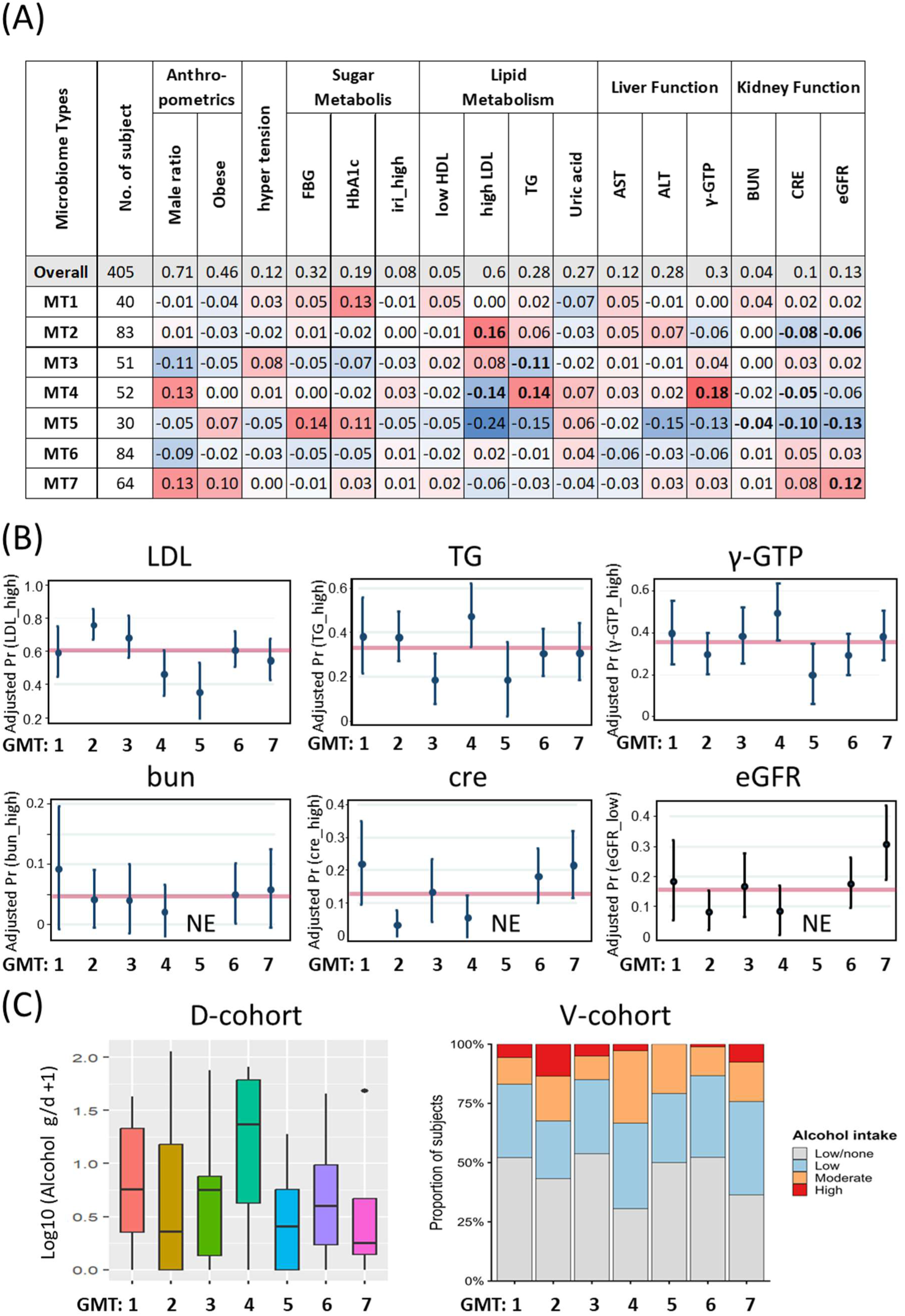
Associations between GMTs and clinical and lifestyle characteristics. **(A)** Heatmap summarizing associations between MT assignment and host clinical parameters estimated by multinomial logistic regression. Clinical variables were standardized by z-score transformation before analysis, and models were adjusted for age, sex, and medication use. Red and blue indicate positive and negative regression coefficients, respectively. Bold values indicate statistically significant associations after multiple testing correction. **(B)** Adjusted differences in the probability of abnormal clinical values for representative clinical parameters across MTs. Probabilities were estimated from logistic regression models adjusted for age, sex, and medication use. Points indicate the difference between the MT-specific adjusted probability and the overall cohort proportion, and error bars represent 95% confidence intervals. The horizontal red line indicates the overall cohort proportion (difference = 0). “NE” indicates that no participants met the predefined abnormality criterion. **(C)** The left panel shows the distribution of daily alcohol intake (log-transformed, g/day) across MTs in the discovery cohort, whereas the right panel shows the distribution of alcohol intake categories across MTs in the validation cohort. Alcohol intake was categorized as Low/none (<23 g ethanol/day), Low (23 to <46 g/day), Moderate (46 to <69 g/day), and High (≥69 g/day), according to the self-administered health check-up questionnaire.

Several lipid and liver function parameters showed distinct GMT-specific patterns. LDL cholesterol was significantly elevated in MT-2, whereas significantly lower levels were observed in MT-4 and MT-5. Triglyceride levels were significantly reduced in MT-3 and elevated in MT-4. MT-4 also exhibited significantly elevated γ-glutamyl transpeptidase (γ-GTP) levels. This was consistent with the higher alcohol consumption observed for MT-4 in both the D- and V-cohorts (Figure 8C), suggesting an association with alcohol-related liver dysfunction. Renal function markers likewise differed among MTs. MT-5 was characterized by lower blood urea nitrogen (BUN) and creatinine levels together with higher estimated glomerular filtration rate (eGFR), consistent with a relatively favorable renal function profile.

No significant GMT differences were observed for glucose-related parameters. Nevertheless, MT-5 tended to exhibit slightly higher glucose-related parameters, a trend that was also apparent in the D-cohort and broadly consistent with its higher BMI profile, although these differences did not reach statistical significance.

Together, these findings suggest that MTs are associated with variation in host physiological parameters, supporting the potential clinical relevance of GMT classification.

### 3.12. Validation of CAG-based microbiome stratification

To further evaluate the transferability of the proposed microbiome stratification framework, we compared the transferred CAG-based GMT classification with de novo genus-based GMT classification constructed independently in the V-cohort (Supplementary Table S28). For comparison, genus-based GMTs were generated by de novo clustering of 229 genus-level abundance profiles from the V-cohort and evaluated using the same multinomial logistic regression framework. De novo genus-based clustering showed explanatory performance broadly comparable to that of the transferred CAG-based GMT classification (McFadden’s pseudo-R² = 0.097 vs. 0.092; Nagelkerke’s R² = 0.296 vs. 0.298), despite the transferred framework requiring neither cohort-specific model reconstruction nor optimization and relying on substantially fewer features. These findings support the transferability of the CAG-based framework across independent cohorts and its potential as a generalizable basis for microbiome stratification.

Finally, we compared the explanatory value of GMTs with that of representative genera characterizing each GMT for their corresponding clinical phenotypes using Firth logistic regression models (Supplementary Tables S29 and S30). MT-4 and MT-5 were selected because they exhibited the strongest and most reproducible associations with liver- and renal-related clinical parameters, respectively. *Faecalibacterium* and *Bifidobacterium* were selected as representative genera of MT-4 and MT-5, respectively, because they consistently characterized these GMTs in both the D- and V-cohorts. When GMTs and their corresponding representative genera were evaluated together, MT-4 and MT-5 retained significant associations with their respective clinical phenotypes, whereas neither *Faecalibacterium* nor *Bifidobacterium* provided additional explanatory value (Supplementary Table S29). These findings indicate that the clinical phenotypes associated with MT-4 and MT-5 were better captured by the community-level GMTs than by their representative genera alone.

To further examine this relationship, we focused on the association between MT-4 and elevated γ-GTP (Supplementary Table S30). Addition of MT-4 to the model significantly improved the explanation of elevated γ-GTP (P = 0.015), whereas addition of *Faecalibacterium* abundance did not (P = 0.210). Moreover, adding *Faecalibacterium* abundance to a model already containing MT-4 did not improve model fit (P = 0.836), whereas adding MT-4 to a model containing *Faecalibacterium* abundance provided significant additional explanatory value (P = 0.037). These findings indicate that the association between MT-4 and elevated γ-GTP was not explained solely by depletion of *Faecalibacterium*.

## 4. Discussion

In the D-cohort, we confirmed that both microbiome composition and metabolite profiles remained relatively stable within individuals over the one-month sampling interval despite substantial inter-individual heterogeneity. These observations motivated the development of a microbial guild-based framework to capture reproducible ecological organization underlying this structural and functional complexity. The 27-CAG-based, seven-GMT framework established in the D-cohort was successfully applied to the V-cohort without redefining either the CAGs or GMTs. The resulting GMT assignments captured variation in gut microbiome organization associated with host clinical characteristics, supporting the validity of this framework as a biologically meaningful representation of microbiome variation.

The 27 CAGs represented more than 70% of the total microbial abundance in both the D- and V-cohorts, indicating that the major ecological structure of the gut microbiome can be represented by a limited number of ecologically coherent co-abundance modules. Furthermore, the observed stability of these CAGs across repeated clustering analyses suggests that these co-abundance patterns reflect underlying ecological organization rather than random variation. The presence of both compact small modules and more loosely connected larger modules further indicates that the gut microbiome is organized across multiple ecological scales, ranging from tightly interacting microbial groups to broader community-level structures. Together, these observations suggest that much of the apparent complexity of the gut microbiome can be represented by a limited number of reproducible ecological modules (CAGs), consistent with previous network-based analyses across geographically distinct human cohorts (Jackson et al., 2018; Loftus et al., 2021). Unlike conventional taxonomy-based classifications, these ecological modules are defined by coordinated co-abundance patterns spanning multiple taxonomic groups. For example, individual taxa such as *Bacteroides* were distributed across multiple CAGs in distinct compositional contexts, indicating that microbial community structure is shaped by community context rather than taxonomic identity alone. These ecological characteristics are consistent with the concept of microbial guilds, representing coherent functional units that capture microbial relationships beyond taxonomic boundaries (Wu et al., 2021).

CAG-based representations showed significant associations with both SCFA and BA profiles, whereas genus-level profiles failed to capture associations with BA profiles. This contrast may reflect fundamental differences in the ecological organization of these microbial metabolic processes. SCFA production is often driven by a limited number of specialized producers (Oliver et al., 2024), allowing taxonomic composition to capture much of the relevant functional variation. In contrast, BA metabolism represents a dynamic host-microbiome process in which host BA secretion is dynamically regulated by diet and host physiological state, thereby reshaping the intestinal BA environment while microbial communities simultaneously modify BA composition (Ridlon et al., 2016; Aoi et al., 2024; Just et al., 2018; Watanabe et al., 2025). Consistent with this dynamic nature of this host-microbiome interaction, BA profiles exhibited substantially greater temporal variability than overall microbiome composition, yet remained significantly associated with CAG organization. These findings suggest that microbial guilds capture coordinated ecological responses to a changing BA environment, highlighting their potential to represent dynamic ecological adaptation to changing BA conditions. A similar ecological perspective may apply to protein fermentation. Unlike the major SCFAs, which primarily reflect carbohydrate fermentation by specialized microbial producers, BCFAs arise from community-level branched-chain amino acid metabolism. The consistent associations of two independent Christensenellaceae-enriched CAGs with multiple BCFAs suggest that these ecological modules may capture community-level protein fermentation activities that are not readily attributable to Christensenellaceae alone.

Previous studies have suggested that gut microbiome composition is better represented as a continuum than as a collection of discrete community types (Knights et al., 2014; Tap et al., 2023). Consistent with this view, our GMT classification likewise showed gradual transitions among microbiome configurations rather than sharply separated clusters. Within this continuous ecological landscape, the seven-GMT framework provided a practical balance between ecological resolution, reproducibility, and biological interpretability. Importantly, despite the absence of clearly defined boundaries, GMTs retained meaningful associations with microbial metabolites and host clinical characteristics when transferred to the independent V-cohort. These findings support the view that GMTs should be regarded not as fixed natural categories but as representative ecological configurations that provide an interpretable framework for describing major patterns of gut microbiome organization.

The identified GMTs exhibited moderate temporal stability, with more than 70% of participants retaining the same microbiome type over the one-month interval, consistent with previous longitudinal studies demonstrating the relative stability of the adult gut microbiome over periods of weeks to months (David et al., 2014; Schlomann and Parthasarathy, 2019). Importantly, GMT transitions were not randomly distributed but occurred preferentially between a limited number of related GMTs, supporting the view that GMTs represent interconnected ecological configurations occupying a constrained ecological state space (Lozupone et al., 2012).

Previous community-based network analyses have shown that ecological organization within the gut microbiome is conserved across independent human cohorts (Jackson et al., 2018; Loftus et al., 2021). Nevertheless, translating microbiome-based classification frameworks across independent cohorts remains challenging, as taxonomic signatures identified in one cohort often exhibit limited generalizability in external datasets (Li et al., 2023). Against this background, the CAG-based GMT framework was successfully transferred to the V-cohort without redefining either CAGs or GMTs, achieving performance comparable to that of de novo genus-level clustering despite requiring neither cohort-specific re-optimization nor de novo model construction. Our findings suggest that CAG-based representations preserve aspects of microbial ecological organization that remain transferable across independent cohorts, thereby preserving microbiome-associated functional and host-associated patterns. Notably, BA-associated functional profiles were more consistently reproduced across cohorts, whereas SCFA-associated profiles retained the major GMT-specific features despite less consistent overall patterns.

Among GMTs, MT-4 exhibited the most distinctive combination of microbial and host-associated features. MT-4 displayed a dysbiosis-like profile characterized by markedly reduced microbial diversity, decreased butyrate levels, and elevated primary BA levels. Together, these features suggest impairment of complementary microbial metabolic functions within the gut ecosystem. MT-4 was also enriched in inflammation-associated taxa, including *Fusobacterium* and *Ruminococcus gnavus*. (Sheng et al., 2022) These microbial features were accompanied by a tendency toward higher alcohol consumption in both cohorts, together with elevated γ-GTP and triglyceride levels, suggesting an association with disturbances in liver-related metabolic function. Interestingly, although depletion of *Faecalibacterium* represented one of the hallmark microbial features of MT-4, the MT-4 classification provided greater explanatory power for the associated clinical phenotype than *Faecalibacterium* abundance alone (Supplementary Table S30). These findings suggest that the clinical characteristics associated with MT-4 emerge from the coordinated organization of the microbial community rather than from the abundance of a single bacterial taxon. The coexistence of reduced butyrate levels, altered bile acid profiles, increased inflammation-associated taxa, and reduced microbial diversity further supports the view that MT-4 represents an ecosystem-level microbiome configuration rather than the expansion or depletion of any individual microbial lineage.

In contrast, MT-5, characterized by a *Bifidobacterium*-enriched microbial guild, was associated with favorable renal function together with favorable lipid-related metabolic characteristics. Interestingly, although *Bifidobacterium* represented a hallmark microbial feature of MT-5, the MT-5 classification showed a stronger association with favorable renal function than *Bifidobacterium* abundance alone (Supplementary Table S29). These findings support the view that the host phenotype associated with MT-5 reflects a guild-level microbial configuration rather than the sole contribution of *Bifidobacterium*. Bifidobacterium has been implicated in the modulation of gut-derived uremic toxins and has been linked to kidney health in previous studies (Rossi et al., 2016). Accordingly, although enrichment of *Bifidobacterium* may contribute to these favorable renal characteristics, the observed host phenotype is more likely to reflect the coordinated ecological organization of the MT-5 microbial guild.

Taken together, this study demonstrates that the ASV-CAG-based approach provides a generalizable framework for microbiome typing that captures functionally relevant variation associated with host physiology across independent cohorts, with potential applicability to diverse human populations. Nevertheless, further optimization of analytical methods, validation in larger and more diverse cohorts, incorporation of longer-term longitudinal designs, and deeper characterization of host phenotypes and microbial functions will be important for establishing the robustness and broader applicability of this framework. Despite these limitations, our findings support microbial guilds as biologically meaningful ecological units and demonstrate that microbiome types defined by combinations of these guilds provide an interpretable ecological framework for understanding functionally relevant variation in the human gut microbiome and its relationship with host physiology.

## Author contributions

J.N.: Conceptualization, Methodology, Software, Validation, Formal analysis, Investigation, Data curation, Writing - original draft, Writing - review & editing, Visualization, Supervision, Project administration, and Funding acquisition. M.Y., A.U., M.T., S.T., and A.S.: Investigation. K.K.: Project administration and Funding acquisition. T.I.: Data curation. H.W.: Investigation, Resources and Data curation. Y.T.: Project administration and Funding acquisition.

## Use of generative AI

Generative artificial intelligence (ChatGPT, OpenAI) was used as a coding assistant to support the development, modification, and debugging of scripts used in some of the analyses, and to assist with English-language editing. All AI-assisted code was reviewed and executed by the authors, and the resulting outputs were verified by the authors. All scientific analyses, interpretations, and conclusions remain the responsibility of the authors.

## Funding

This work was supported in part by Grants-in-Aid for Scientific Research from the Japan Society for the Promotion of Science (JSPS KAKENHI; Grant Nos. 17H04620, 20KK0130, and 22K19136 to J.N.; Grant Nos. 15K09015 and 22K08037 to T.I.), the JSPS Core-to-Core Program “Establishment of Gut Microbiome Research Core Linking Asian Foods and Health,” the Kyushu University Institute for Asian and Oceanian Studies, grants from the Mishima Kaiun Memorial Foundation and the Kieikai Research Foundation, and research funding from Ajinomoto Co., Inc.

## Acknowledgements

We sincerely thank all study participants for their participation and for providing biological samples and clinical information. We also thank the staff of Tohoku Central Hospital for their assistance with participant recruitment, sample collection, and clinical data collection. We appreciate the technical assistance with the operation of the MiSeq and LCMS-8050 provided by the Center for Advanced Instrumental and Educational Supports, Faculty of Agriculture, Kyushu University.

## Data availability

The 16S rRNA gene amplicon sequencing data generated in this study have been deposited in the DDBJ Sequence Read Archive under BioProject accession number PRJDB5860. The sequencing data for the D- and V-cohorts are available under Run accession numbers DRR1087786-DRR1088017 and DRR1088018-DRR1088421, respectively.

## Competing interests

This study was partially supported by research funding from Ajinomoto Co., Inc. The authors declare no other competing interests.

## Abbreviations

ASV: amplicon sequence variant
CAG: co-abundance group
GMT: gut microbiome type
CLR: centered log-ratio transformation
PERMANOVA: permutational multivariate analysis of variance
SCFA: short-chain fatty acid
BA: bile acid
CA: cholic acid
CDCA: chenodeoxycholic acid
DCA: deoxycholic acid
LCA: lithocholic acid
UDCA: ursodeoxycholic acid
GCA: glycocholic acid
GCDCA: glycochenodeoxycholic acid
GDCA: glycodeoxycholic acid
GLCA: glycolithocholic acid
GUDCA: glycoursodeoxycholic acid
TCA: taurocholic acid
TCDCA: taurochenodeoxycholic acid
TDCA: taurodeoxycholic acid
TLCA: taurolithocholic acid
TUDCA: tauroursodeoxycholic acid

