## Supplementary Note for "Seven gut microbiome types across two Japanese cohorts defined by co-abundance-based microbial guilds and associated with host physiology"

### Supplementary Materials

#### Supplementary Note 1. Study design and cohorts

This study was conducted using two independent Japanese cohorts: a discovery cohort and an external validation cohort.

In the discovery cohort, participants were instructed to defer stool collection if they had experienced influenza or acute gastroenteritis within the preceding two weeks or were currently symptomatic. Stool samples were collected only after resolution of diarrheal symptoms. Participants were also asked to refrain from excessive alcohol consumption, intense physical activity, and consumption of spicy or otherwise irritative foods on the day before and the day of sampling. These instructions were intended to reduce transient influences on gut microbiome composition.

Participants in the discovery cohort were recruited through personal networks of the research team, including former laboratory members and their acquaintances. The discovery cohort consisted of 116 community-dwelling adults aged 19–76 years living in Japan. Fecal samples were collected twice from each participant at a one-month interval and analyzed for bacterial composition, short-chain fatty acids (SCFAs), and bile acids (BAs).

Between the two sampling time points, dietary intake, including alcohol consumption, was assessed using a validated self-administered diet history questionnaire (DHQ), a long-form questionnaire designed to capture detailed habitual dietary patterns over the preceding month. [41,42] In addition, information on medical history, current medication use, smoking habits (past and current), and bowel conditions was collected via questionnaires. Detailed participant information is provided in Supplementary Table S1 for demographic and clinical variables and in Supplementary Table S2 for dietary data.

The validation cohort consisted of 404 community-dwelling adults aged 42–62 years who underwent routine health check-ups at Tohoku Chuo Hospital. Fecal samples were collected once prior to the health check-up and analyzed for microbiome, SCFA, and BA analyses. Clinical measurements obtained during the health check-ups were used for subsequent association analyses with microbiome types. Compared with the discovery cohort, fewer restrictions were applied to sample collection conditions in the validation cohort, allowing assessment of microbiome variation across individuals with diverse physiological and clinical backgrounds. Detailed participant information for the validation cohort is provided in Supplementary Table S11.

The overall workflow of the study is shown in Fig. 1 together with the corresponding figures and tables. Supplementary Tables S1–S10 correspond to analyses of the discovery cohort, whereas Supplementary Tables S11–S16 correspond to analyses of the validation cohort. Supplementary Tables S1 and S2 provide demographic, clinical, and dietary information for the discovery cohort, while Supplementary Table S11 summarizes clinical measurements obtained during routine health check-ups in the validation cohort.

#### Supplementary Note 2. Construction of co-abundance networks and identification of ASV-based CAGs

##### 1. Overall workflow

The overall workflow for construction and evaluation of ASV-based CAGs is summarized in Supplementary Table S17.

##### 2. ASV filtering

A total of 3,523 ASVs identified from the D-cohort (Supplementary Table S3) were filtered using a standardized workflow. For each sample, an ASV was considered present if its relative abundance was ≥0.05% or if it was supported by at least 10 sequencing reads. Based on this criterion, prevalence across samples was calculated, and ASVs detected in fewer than 1% of the 232 samples were removed. This procedure retained 1,189 ASVs for downstream network analysis.

| Supplementary Table S17. Workflow for construction and evaluation of ASV-based co-abundance groups (CAGs). | | | | |
|---|---|---|---|---|
| **No.** | **Step** | **Method** | **Program** | **Output** |
| 1 | ASV inference | Denoising and ASV inference | QIIME2 (DADA2) | 3,523 ASVs (Table S3) |
| 2 | ASV filtering | Quality and prevalence-based filtering | custom script | 1,189 retained ASVs |
| 3 | Correlation inference | Compositional correlation analysis with bootstrap resampling | FastSpar v1.0.0 | Pairwise ASV correlation matrix |
| 4 | Significance assessment | Multiple-testing correction | Benjamini–Hochberg method | Significant ASV correlations |
| 5 | Co-abundance network construction | Network sparsification and weighted network construction | igraph (R) | Weighted co-abundance network |
| 6 | Community detection | Modularity optimization | Louvain algorithm (igraph) | 27 CAGs with 316 ASVs (Table S4) |
| 7 | Network visualization | Visualization of CAG network | Cytoscape v3.10.3 | Network figure (Figure 3) |
| 8 | Robustness evaluation | Comparison of clustering solutions | R (NMI, ARI, module-size comparison) | Robustness statistics (Table S18) |
| 9 | CAG abundance calculation | Aggregation of ASV abundances within each CAG | Custom R scripts | Sample-by-CAG abundance matrix (Table S6) |
| 10 | Coverage evaluation | Assessment of CAG representation | Custom R scripts | Coverage metrics |

#### 3. FastSpar correlation analysis

Pairwise associations among ASVs were estimated using FastSpar (v1.0.0), [49] with 100 iterations and 10 exclusion iterations. To evaluate statistical robustness, 1,000 bootstrap datasets were generated using fastspar bootstrap, and pairwise correlations were re-estimated independently for each bootstrap dataset using FastSpar. Empirical p-values were calculated as the proportion of bootstrap correlations exceeding the observed correlation and were adjusted using the Benjamini–Hochberg procedure[51] to obtain q-values. Association stability was further evaluated using two reproducibility metrics. Selection frequency was defined as the proportion of bootstrap replicates in which an edge satisfied the predefined significance criteria, whereas sign consistency represented the proportion of bootstrap replicates preserving the direction of the inferred correlation.

#### 4. Network construction

Edges satisfying $r \geq 0.20$ and $q \leq 0.10$ were retained. To reduce spurious associations while maintaining network connectivity, a k-nearest-neighbor-inspired sparsification procedure was applied by retaining, for each ASV, up to the eight strongest associations ranked by absolute correlation coefficient. The union of retained edges defined the final network.

#### 5. Community detection

An undirected weighted network was constructed using the igraph package, [52] with absolute correlation coefficients used as edge weights and the original correlation sign retained as an edge attribute. Community detection was performed using the Louvain algorithm, [53] and each detected module was defined as a co-abundant group (CAG). Structural properties of each CAG, including node count, number of internal edges, edge density, mean internal edge weight, proportions of positive and negative edges, and cut ratio, were calculated (Supplementary Table S4). Network visualization was performed using Cytoscape v3.10.3 with an Edge-Weighted Spring-Embedded layout. Node size was scaled according to mean relative abundance, and node color represented family-level taxonomic affiliation.

#### 6. Robustness analysis

To evaluate robustness of community detection, Louvain clustering was repeated using five different random seeds. Concordance among clustering solutions was assessed using normalized mutual information (NMI) and adjusted Rand index (ARI) as shown in Supplementary Table S19. In addition, CAG module-size distributions were compared across clustering solutions as an additional measure of clustering robustness (Supplementary Table S20). The clustering solution generated with the initial random seed (1234) was retained for all downstream analyses, whereas additional random seeds were used to evaluate clustering stability.

**Supplementary Table S4. Topological properties of co-abundance groups (CAGs)**

| CAG | nodes | edges | density | mean_weight | cut_ratio | top_genera |
|---|---|---|---|---|---|---|
| 1 | 18 | 24 | 0.157 | 0.289 | 0.200 | *Bacteroides, Megasphaera, Parabacteroides* |
| 2 | 27 | 38 | 0.108 | 0.244 | 0.406 | *Incertae_Sedis, Christensenellaceae_R-7_group, Acutalibacter* |
| 3 | 61 | 309 | 0.169 | 0.284 | 0.144 | *Blautia, Faecalibacterium* |
| 4 | 25 | 29 | 0.097 | 0.271 | 0.482 | *Anaerostipes, Blautia, Bacteroides* |
| 5 | 5 | 5 | 0.500 | 0.224 | 0.000 | *Incertae_Sedis, Faecalibacterium, CAG-352* |
| 6 | 8 | 7 | 0.250 | 0.213 | 0.000 | *Bacteroides, Agathobacter, Alistipes* |
| 7 | 13 | 16 | 0.205 | 0.232 | 0.158 | *Christensenellaceae_R-7_group, Incertae_Sedis, UCG-002* |
| 8 | 33 | 74 | 0.140 | 0.272 | 0.169 | *Bacteroides, Blautia, Enterocloster* |
| 9 | 7 | 6 | 0.286 | 0.225 | 0.000 | *Sutterella, Segatella, Senegalimassilia* |
| 10 | 23 | 26 | 0.103 | 0.257 | 0.278 | *Parabacteroides, [Eubacterium]_eligens_group, Anaerobutyricum* |
| 11 | 3 | 3 | 1.000 | 0.350 | 0.000 | *Streptococcus* |
| 12 | 25 | 32 | 0.107 | 0.255 | 0.000 | *Segatella, Dialister, Mitsuokella* |
| 13 | 2 | 1 | 1.000 | 0.211 | 0.500 | *Bacteroides, Incertae_Sedis* |
| 14 | 5 | 4 | 0.400 | 0.268 | 0.000 | *Bacteroides* |
| 15 | 4 | 3 | 0.500 | 0.325 | 0.400 | *Bacteroides, Megamonas* |
| 16 | 9 | 9 | 0.250 | 0.254 | 0.500 | *Bifidobacterium, Romboutsia, Terrisporobacter* |
| 17 | 14 | 17 | 0.187 | 0.271 | 0.227 | *Bacteroides, Veillonella, Acidaminococcus* |
| 18 | 5 | 4 | 0.400 | 0.377 | 0.000 | *Megasphaera, Bacteroides* |
| 19 | 5 | 4 | 0.400 | 0.276 | 0.000 | *Parabacteroides, Bacteroides* |
| 20 | 4 | 3 | 0.500 | 0.228 | 0.000 | *Bacteroides, Parabacteroides* |
| 21 | 3 | 2 | 0.667 | 0.301 | 0.000 | *Fusobacterium, Megamonas* |
| 22 | 5 | 5 | 0.500 | 0.258 | 0.000 | *Megamonas, Megasphaera, Bacteroides* |
| 23 | 3 | 2 | 0.667 | 0.233 | 0.000 | *Parasutterella, Parabacteroides* |
| 24 | 2 | 1 | 1.000 | 0.277 | 0.667 | *Sutterella* |
| 25 | 3 | 2 | 0.667 | 0.223 | 0.000 | *Flavonifractor, Butyricicoccus, Blautia* |
| 26 | 2 | 1 | 1.000 | 0.219 | 0.000 | *Incertae_Sedis* |
| 27 | 2 | 1 | 1.000 | 0.259 | 0.000 | *Extibacter, Sellimonas* |

Each row represents a CAG identified by network-based clustering.

Modules are characterized by the number of ASVs (nodes), number of edges, network density, number of positive and negative correlations, mean correlation strength, and cut ratio.

Dominant genera are shown based on the most abundant taxa within each module.

**Supplementary Table S19.** Pairwise similarity of CAG assignments obtained with different random seeds, assesssed using the adjusted Rand indes (ARI) and normalized mutual information (NMI).

| | | ARI | | | | NMI | | | |
|---|---|---|---|---|---|---|---|---|---|
| **seed** | **No of Modules** | **1002** | **1003** | **1004** | **1005** | **1002** | **1003** | **1004** | **1005** |
| **1001** | 27 | 0.736 | 0.701 | 0.786 | 0.748 | 0.894 | 0.871 | 0.909 | 0.896 |
| **1002** | 26 | | 0.888 | 0.808 | 0.790 | | 0.938 | 0.907 | 0.912 |
| **1003** | 26 | | | 0.751 | 0.759 | | | 0.873 | 0.889 |
| **1004** | 27 | | | | 0.816 | | | | 0.911 |
| **1234** | 27 | | | | | | | | |
| | | **Overall mean ARI = 0.778** | | | | **Overall mean NMI = 0.900** | | | |

*Seed 1234 correponds to the clustering solution used for all downstream analyses.

**Supplementary Table S20.** Sizes of ranked CAG modules across Louvain clustering runs using different random seeds.

| CAG rank | 1234 | 1001 | 1002 | 1003 | 1004 |
|---|---|---|---|---|---|
| 1 | 55 | 51 | 54 | 51 | 61 |
| 2 | 41 | 44 | 46 | 34 | 33 |
| 3 | 37 | 36 | 38 | 29 | 27 |
| 4 | 25 | 25 | 25 | 26 | 25 |
| 5 | 21 | 25 | 19 | 25 | 25 |
| 6 | 16 | 22 | 17 | 24 | 23 |
| 7 | 14 | 16 | 16 | 21 | 18 |
| 8 | 13 | 13 | 15 | 14 | 14 |
| 9 | 11 | 11 | 13 | 11 | 13 |
| 10 | 10 | 8 | 8 | 8 | 9 |
| 11 | 8 | 7 | 7 | 8 | 8 |
| 12 | 7 | 7 | 7 | 7 | 7 |
| 13 | 7 | 5 | 5 | 7 | 5 |
| 14 | 5 | 5 | 5 | 5 | 5 |
| 15 | 5 | 5 | 5 | 5 | 5 |
| 16 | 5 | 5 | 5 | 5 | 5 |
| 17 | 5 | 5 | 5 | 5 | 5 |
| 18 | 5 | 4 | 4 | 5 | 4 |
| 19 | 4 | 4 | 4 | 4 | 4 |
| 20 | 4 | 3 | 3 | 4 | 3 |
| 21 | 3 | 3 | 3 | 3 | 3 |
| 22 | 3 | 3 | 3 | 3 | 3 |
| 23 | 3 | 3 | 3 | 3 | 3 |
| 24 | 3 | 2 | 2 | 3 | 2 |
| 25 | 2 | 2 | 2 | 2 | 2 |
| 26 | 2 | 2 | 2 | 2 | 2 |
| 27 | 2 | | | 2 | 2 |

CAG modules were ranked by the number of ASVs within each clustering run. The clustering solution generated with random seed 1234 (red bars) was used for downstream analyses.

##### 7. Construction of CAG abundance matrix

For each sample, relative abundances of ASVs belonging to the same CAG were summed to generate a sample-by-CAG abundance matrix (Supplementary Table S6). Relative abundances were normalized by library size before downstream analyses.

To assess representativeness of the resulting CAGs, three coverage metrics were calculated: ASV coverage (proportion of ASVs assigned to CAGs), abundance coverage (mean proportion of total community abundance represented by CAG-associated ASVs), and variance coverage (proportion of total among-sample variance in ASV relative abundances explained by CAG-associated ASVs). The retained CAGs accounted for 8.97% of all ASVs, while representing 75.9% of the total community abundance and 79.5% of the total sample variance.

#### Supplementary Note 3. Association of CAGs with SCFAs and BAs.

For association analyses, SCFA metabolites included acetate, propionate, butyrate, succinate, lactate, isobutyrate, isovalerate, and valerate. BA metabolite groups comprised primary bile acids (CA + CDCA), secondary bile acids (DCA + LCA), UDCA, and conjugated bile acids.

##### 1. Redundancy analysis (RDA)

RDA was performed using the rda() function in the vegan package (version 2.7.2) in R (version 4.1.2). Relative abundances of CAGs were used as explanatory variables, whereas measured fecal concentrations of eight SCFAs and four metabolically defined BA groups were used as response variables without compositional or relative-abundance transformation. Before RDA, both the explanatory and response variables were centered and scaled to unit variance. Samples containing missing values for any response variable were excluded from the corresponding analysis. Overall model significance was evaluated using permutation tests with 999 permutations. The significance of individual CAG terms was further assessed using sequential term-wise permutation tests implemented with anova.cca() (by = "terms"). Adjusted coefficients of determination were calculated using RsquareAdj(). The result of RDA is summarized in Table 2.

**2. Correlation analysis between CAG and SCFA/BA**

Pairwise correlations between individual CAGs relative abundances and concentrations of each SCFA and BA metabolite group were evaluated using Spearman's rank correlation analysis in Stata/SE 12.0 (StataCorp LP, College Station, TX, USA). The correlation results are visualized in a Sankey diagram (Fig. 4) and a heatmap (Fig. 5). Sankey diagrams were generated to visualize the hierarchical relationships among bacterial genera, CAGs, SCFAs, and BA groups. The Sankey diagram was generated using the networkD3 package in R and exported as an editable SVG graphic. Links between CAGs and metabolites represent statistically significant associations ($P < 0.05$). Each CAG was assigned to the dominant bacterial genus according to the taxonomic annotation of its constituent ASVs. Links between genera and CAGs represent taxonomic membership, whereas links between CAGs and metabolites represent statistically significant associations identified by Spearman's rank correlation analysis. Only statistically significant associations were included to emphasize the hierarchical links between bacterial taxonomic composition, CAGs, and metabolite profiles.

#### Supplementary Note 4. CAG-based microbiome typing and validation.

##### 1. Construction of CAG-based microbiome typing (GMTs)

The relative abundances of CAGs in each sample were re-normalized to sum to one before distance calculation. Jensen–Shannon divergence (JSD) [55] was calculated from the resulting relative abundance profiles, and partitioning around medoids (PAM) clustering [56] was used to identify gut microbiome types (GMTs). For comparison, hierarchical clustering using Ward's method based on Aitchison distance[57] was also evaluated after centered log-ratio (CLR) transformation using a pseudocount of 0.5.

Candidate numbers of clusters (K = 3–10) were systematically evaluated. The final GMT solution was selected by jointly considering average silhouette width, cluster size, and the explanatory power of each clustering solution for fecal SCFA and BA profiles (Supplementary Figure S1 and Supplementary Table S21). Explanatory power was assessed as the adjusted $R^2$ of separate redundancy analyses (RDAs), using GMT assignment as the explanatory variable and either fecal SCFA or BA profiles as the response variables.

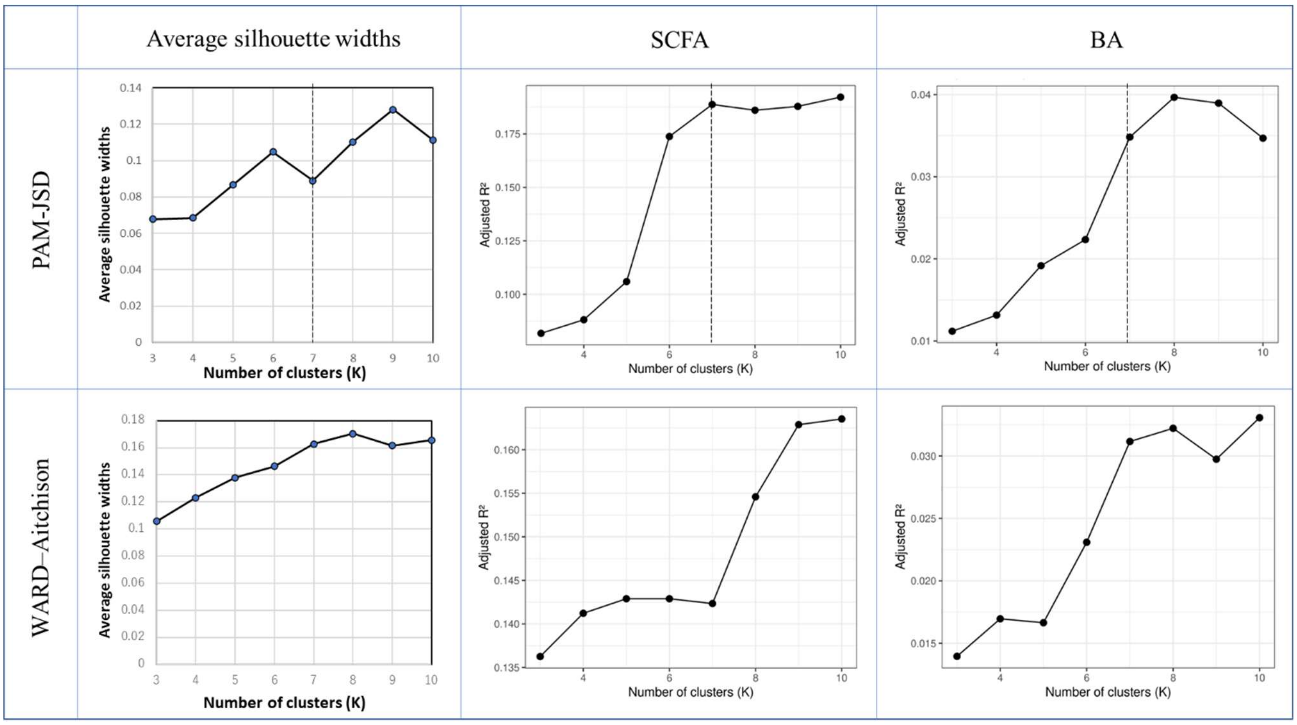


**Supplementary Figure S1. Evaluation of candidate clustering solutions for gut microbiome typing.** Candidate cluster numbers (K = 3–10) were evaluated using mean silhouette width (left), adjusted $R^2$ from redundancy analysis (RDA) based on fecal short-chain fatty acid (SCFA) profiles center), and adjusted $R^2$ from RDA based on bile acid (BA) profiles (right). The upper panels show results obtained using partitioning around medoids (PAM) clustering based on Jensen–Shannon divergence, whereas the lower panels show results obtained using Ward hierarchical clustering based on Aitchison distance. Dashed vertical lines indicate the selected seven-cluster solution.

| PAM_JSD | K=3 | K=4 | K=5 | K=6 | K=7 | K=8 | K=9 | K=10 |
|---|---|---|---|---|---|---|---|---|
| **MT-1** | 103 | 96 | 86 | 86 | 50 | 41 | 41 | 35 |
| **MT-2** | 62 | 61 | 54 | 40 | 39 | 37 | 34 | 6 |
| **MT-3** | 67 | 61 | 53 | 52 | 32 | 32 | 32 | 33 |
| **MT-4** | | 14 | 25 | 15 | 15 | 12 | 12 | 28 |
| **MT-5** | | | 14 | 25 | 25 | 15 | 15 | 12 |
| **MT-6** | | | | 14 | 57 | 25 | 22 | 22 |
| **MT-7** | | | | | 14 | 56 | 56 | 15 |
| **MT-8** | | | | | | 14 | 14 | 21 |
| **MT-9** | | | | | | | 6 | 48 |
| **MT-10** | | | | | | | | 12 |

| WARD_AIT | K=3 | K=4 | K=5 | K=6 | K=7 | K=8 | K=9 | K=10 |
|---|---|---|---|---|---|---|---|---|
| **MT-1** | 89 | 89 | 89 | 89 | 89 | 89 | 89 | 89 |
| **MT-2** | 123 | 114 | 34 | 34 | 34 | 34 | 34 | 9 |
| **MT-3** | 20 | 20 | 80 | 80 | 70 | 58 | 14 | 34 |
| **MT-4** | | 9 | 20 | 8 | 10 | 10 | 44 | 3 |
| **MT-5** | | | 9 | 12 | 8 | 8 | 10 | 44 |
| **MT-6** | | | | 9 | 12 | 12 | 8 | 10 |
| **MT-7** | | | | | 9 | 12 | 12 | 8 |
| **MT-8** | | | | | | 9 | 12 | 11 |
| **MT-9** | | | | | | | 9 | 12 |
| **MT-10** | | | | | | | | 12 |

**Supplementary Table S21.** Cluster size distributions for candidate gut microbiome type (GMT) solutions across different numbers of clusters (K = 3–10). The left and right tables present the results of PAM–JSD and Ward–Aitchison clustering, respectively.

#### 2. Validation of GMT classification

Cluster separation was evaluated by principal coordinate analysis (PCoA) and tested using permutational multivariate analysis of variance (PERMANOVA) with 999 permutations. Homogeneity of multivariate dispersion was assessed using betadisper analysis (Supplementary Figure S2).

Clustering stability was evaluated using an 80% subsampling bootstrap procedure repeated 100 times. Agreement between bootstrap-derived and original clustering solutions was quantified using the adjusted Rand index (ARI) and normalized mutual information (NMI). Reproducibility of GMT classification was assessed using nearest-centroid classification with 10-fold cross-validation (Supplementary Figure S3).

To visualize the CAG configurations underlying the identified gut microbiome types (GMTs), the mean relative abundance of each CAG was calculated separately for each GMT. Mean CAG abundances were standardized across GMTs using Z-score transformation to facilitate comparison of enrichment patterns among GMTs. The resulting Z-score matrix was visualized as a heatmap, and CAGs were hierarchically clustered based on their standardized abundance profiles (Supplementary Figure S4). To facilitate biological interpretation of the identified CAGs, the taxonomic composition of each CAG was displayed as dot plots of its constituent ASVs on the right side of the heatmap.

Supplementary Figure S4 shows that several CAGs exhibited preferential enrichment in specific GMTs, indicating that each microbiome type was characterized by a unique configuration of co-abundance structures rather than by enrichment of a single dominant module. These CAG configurations were accompanied by distinct genus-level compositions (Figure 5A). For example, MT-4 was characterized by enrichment of Bacteroidaceae-dominant CAG1 and CAG14 together with Fusobacteriaceae-dominant CAG21, whereas MT-6 was enriched in Ruminococcaceae-dominant CAG5, CAG7, and CAG26. MT-5 and MT-7 were distinguished by selective enrichment of Bifidobacteriaceae-dominant CAG16 and Prevotellaceae-dominant CAG12, respectively.

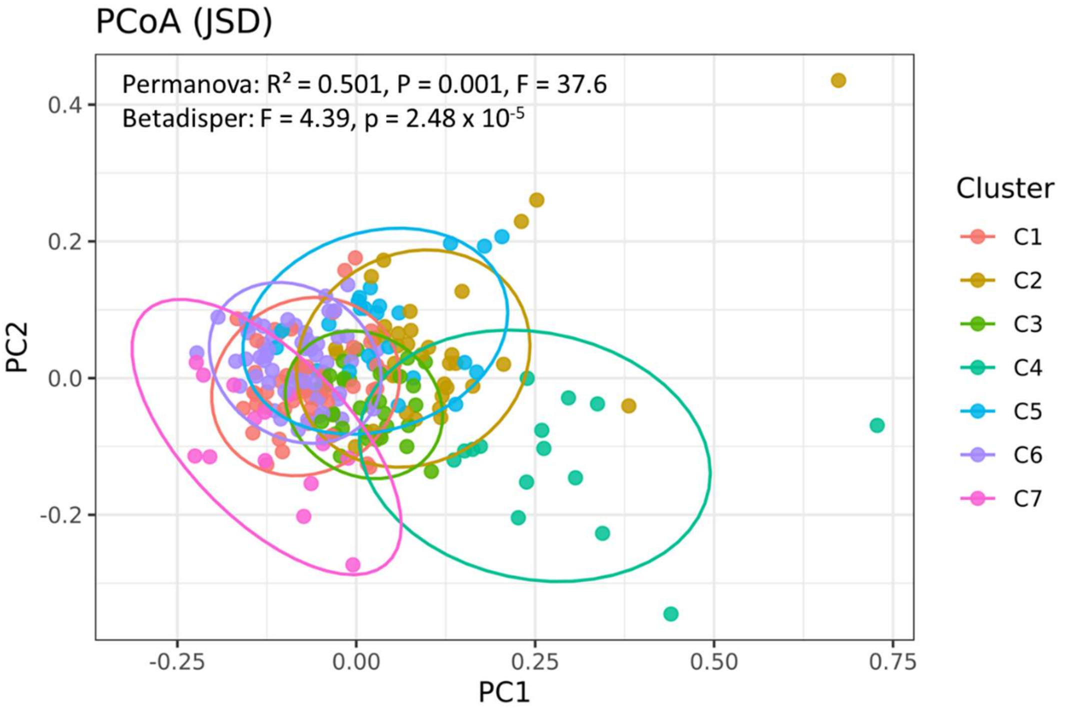


**Supplementary Figure S2. Principal coordinate analysis (PCoA) of discovery cohort samples based on Jensen−Shannon divergence of CAG profiles.** Samples are colored according to GMT assignment obtained by PAM-JSD clustering (k = 7). Ellipses represent 95% confidence intervals for each cluster. PERMANOVA and betadisper statistics are shown to indicate differences in community structure and dispersion among GMTs.

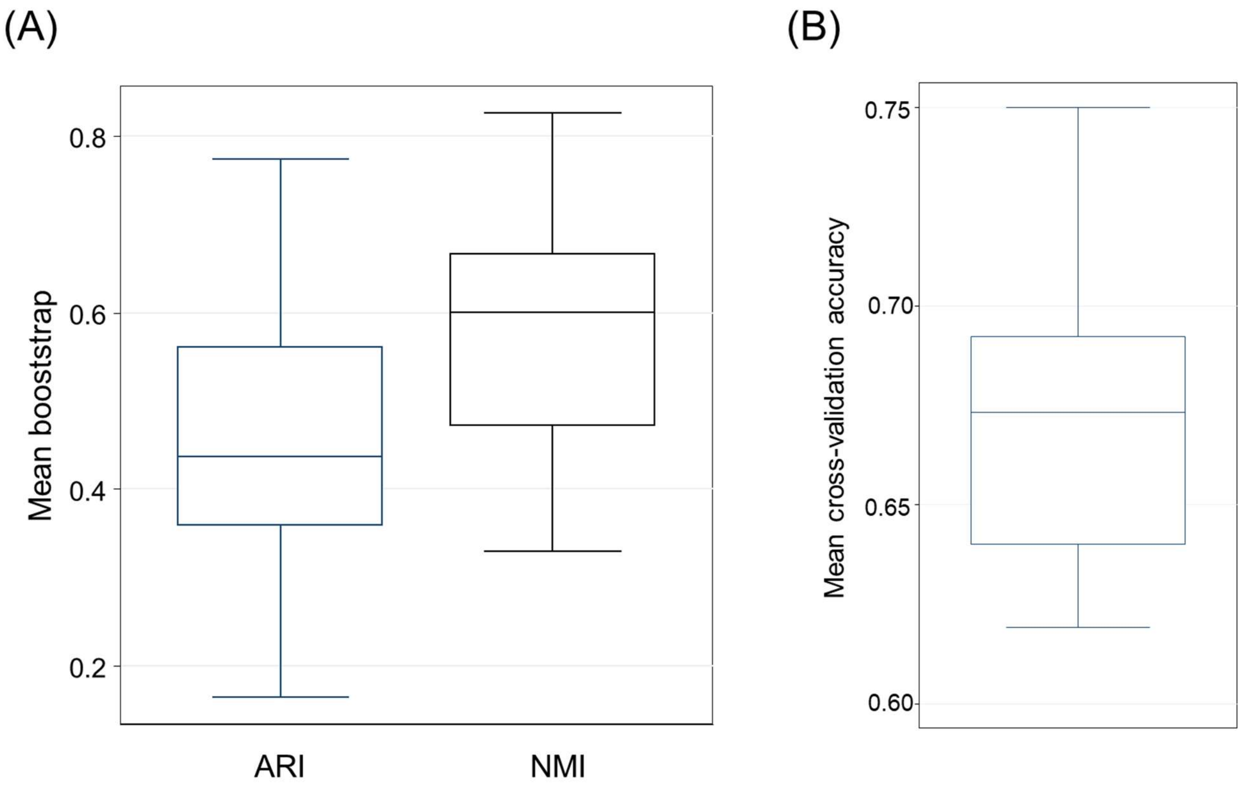


**Supplementary Figure S3.** Robustness and reproducibility of GMT clustering. (A) Distribution of adjusted Rand index (ARI) and normalized mutual information (NMI) values across 100 subsampling-based bootstrap iterations using 80% of samples, demonstrating reproducibility of the clustering structure across resampled datasets. (B) Distribution of classification accuracies obtained by 10-fold cross-validation of GMT assignment. Boxes indicate the interquartile range (IQR), center lines indicate medians, whiskers represent 1.5 × IQR, and points indicate outliers.

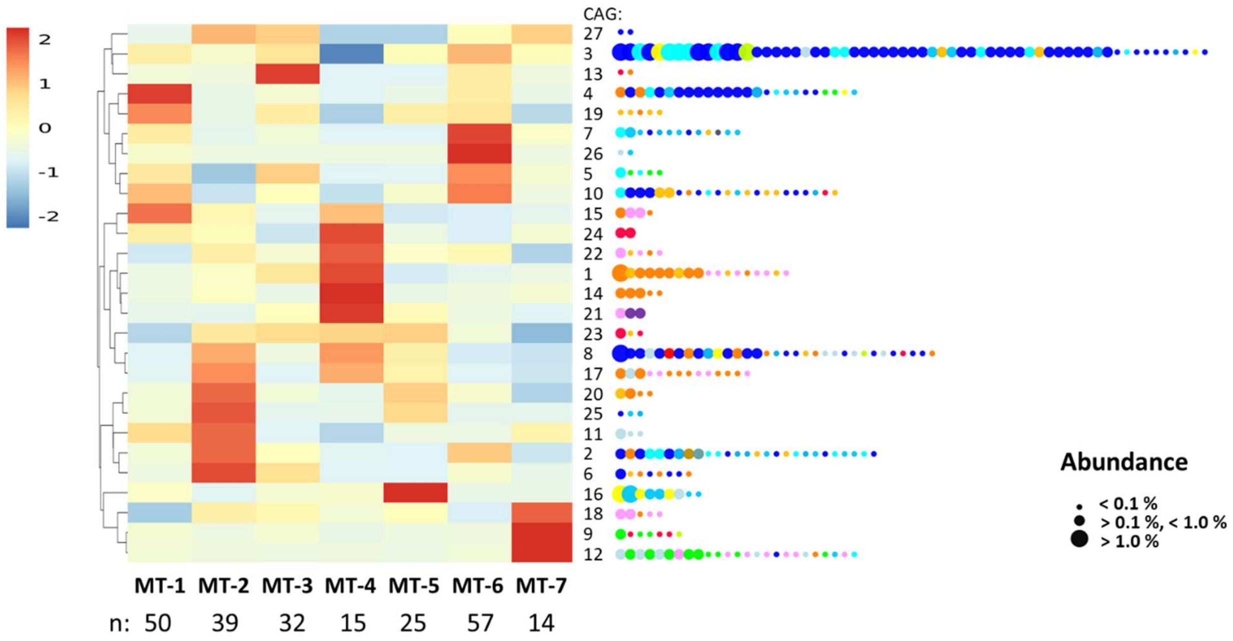


**Supplementary Figure S4. CAG configurations underlying the seven gut microbiome types (GMTs).** A total of 232 samples were classified into seven gut microbiome types (GMTs) based on the relative abundance profiles of 27 CAGs. The left panel shows the mean relative abundance of each CAG across GMTs as a heatmap. Mean CAG abundances were standardized by Z-score across GMTs to highlight relative enrichment or depletion, and CAGs were hierarchically clustered according to their standardized abundance profiles. The heatmap color scale represents Z-score-standardized relative abundances. The right panel shows the taxonomic composition of each CAG as dot plots of its constituent ASVs, where dot size represents relative abundance and colors indicate bacterial families as shown in Fig. 3A.

#### Supplementary Note 5. Characterization of GMT-associated microbial and metabolic features.

Overall differences in fecal SCFA and BA profiles among GMTs were evaluated separately using permutational multivariate analysis of variance (PERMANOVA). Associations between individual GMTs and genus-level microbial composition (Supplementary Table S9), alpha diversity indices, individual SCFAs, and BAs (Supplementary Table S10) were subsequently assessed using MaAsLin2. [66]

Differential analyses of genus-level composition, alpha diversity indices, SCFAs, and BAs were performed using MaAsLin2. For each GMT, a binary indicator variable (cluster_is_k) representing membership in the target GMT was generated, and one-versus-rest comparisons were conducted across all samples using linear models (expr ~ cluster_is_k). No additional normalization or transformation was applied. For genus-level analyses, low-prevalence taxa were excluded using the default abundance and prevalence filters implemented in MaAsLin2, whereas all alpha diversity indices, SCFAs, and BAs were retained. P values were adjusted for multiple testing using the Benjamini–Hochberg false discovery rate (FDR) procedure. Genus-level differential abundance analyses were performed after filtering low-prevalence taxa. Only genera detected in at least 10% of samples (≥24 of 232 samples) were retained, resulting in 129 genera for downstream analyses.

#### Supplementary Note 6. Assessment and visualization of temporal GMT stability.

Transition counts were summarized for all T0–T1 GMT combinations. GMT-specific stability was defined as the proportion of subjects assigned to a given GMT at T0 who retained the same GMT assignment at T1. GMT assignments for all subjects are provided in Supplementary Table S7. Transition probabilities were calculated by dividing the number of subjects undergoing each T0–T1 transition by the total number of subjects assigned to the corresponding GMT at T0, resulting in a row-normalized transition matrix. Subject-level GMT transitions were visualized using a Sankey diagram generated with the ggalluvial package in R (Figure 6A). Flow widths were proportional to the number of subjects undergoing each transition. The transition probability matrix was visualized as a heatmap (Figure 6B). No inferential statistical tests were performed because the analysis was intended to descriptively summarize temporal stability and transition patterns.

#### Supplementary Note 7. Association analyses of GMTs with host-related variables.

Information on host-related variables was obtained using self-administered questionnaires completed by the participants. The variables included in each analysis are summarized in Supplementary Table S1 (host-related variables) and Supplementary Table S2 (dietary variables). Variables were grouped into the following categories and analyzed separately:

**Physiological characteristics:** age, sex, and body mass index (BMI).

**Lifestyle factors:** smoking status, physical activity level (Sport_PAL), alcohol consumption, and total energy intake.

**Disease history:** past medical history variables.

**Infection history:** viral enteritis, influenza, influenza without vaccination, influenza after vaccination, bacteriogenic enteritis, and other infections.

**Medication and gastrointestinal characteristics:** probiotic use, antibiotic use within the previous month, stool frequency, and Bristol stool form scale.

**Dietary variables:** food groups, macronutrients (protein, fat, carbohydrate, and dietary fiber), fatty acid intake (total fatty acids, saturated, monounsaturated, and polyunsaturated fatty acids), and micronutrients. Dietary intake was assessed using the self-administered Diet History Questionnaire (DHQ), which estimates habitual food intake. Estimated food intake was subsequently converted into nutrient intake values using the corresponding nutrient composition database. [41,42]

For each variable group, the same MaAsLin2 analytical framework was applied. For each GMT, one-versus-rest analyses were performed using GMT membership (target GMT versus all remaining GMTs) as the sole fixed effect (expr ~ cluster_is_k). Continuous variables were standardized by z-score transformation, whereas no normalization or data transformation was applied. The results are presented in Supplementary Table S22.

**Supplementary Table S22.** Associations of host-related variables with gut microbiome types (GMTs) estimated by cluster-wise MaAsLin2 analyses.

| Category | feature | Regression coefficient (β) (bold: q < 0.1) | | | | | | |
|---|---|---|---|---|---|---|---|---|
| | | MT-1 | MT-2 | MT-3 | MT-4 | MT-5 | MT-6 | MT-7 |
| Host physiology | Age (y) | 0.113 | 2.654 | -1.359 | 0.613 | -1.364 | -0.587 | -0.573 |
| Host physiology | Sex (male:1/female:2) | -0.112 | **0.159** | 0.001 | -0.065 | -0.007 | 0.013 | 0.001 |
| Host physiology | BMI (kg/m$^2$) | 0.035 | -0.615 | -0.476 | 0.788 | 0.554 | 0.158 | -0.241 |
| Lifestyle factors | Alcohol (g/d) | -1.441 | 2.799 | -0.536 | **6.070** | -2.844 | -2.112 | -0.554 |
| Lifestyle factors | antibiotics in 1 month (no:0/yes:1) | -0.007 | -0.006 | -0.004 | -0.003 | -0.004 | 0.022 | -0.003 |
| Lifestyle factors | Smoking (no:0/yes:1) | -0.013 | 0.029 | -0.027 | **0.073** | 0.017 | -0.040 | -0.018 |
| Lifestyle factors | Sport_PAL | **0.158** | -0.042 | -0.045 | 0.022 | 0.026 | -0.087 | -0.056 |
| Lifestyle factors | Energy Intake (kcal/d) | ##### | 12.339 | ##### | 86.686 | 75.773 | ##### | 33.325 |
| Gut-related factors | Bristol score | 0.087 | -0.195 | 0.169 | -0.069 | 0.083 | -0.034 | -0.037 |
| Gut-related factors | Probiotics (no:0/yes:1) | 0.001 | -0.093 | 0.038 | -0.065 | -0.048 | **0.128** | 0.001 |
| Gut-related factors | Stool frequency (times per week) | -0.200 | 0.229 | -0.022 | -0.094 | -0.101 | 0.199 | -0.104 |

Associations of host physiological, lifestyle, and gut-related factors with GMTs estimated by clusterwise MaAsLin2 analysis. Values represent MaAsLin2 regression coefficients (effect sizes) from one-versus-rest analyses for each GMT. Positive coeffcients indicate enrichment of the corresponding host variable in the indicated GMT, whereas negative coefficients indicate depletion relative to the remaining GMTs.

#### Supplementary Note 8. Transfer of the D-cohort CAG–GMT framework to the validation cohort.

To evaluate the transferability of the CAG-based classification framework, samples in the validation cohort were assigned to the predefined GMTs established in the discovery cohort without re-clustering or model re-training.

**1. Transfer of predefined CAGs from D-cohort to V-cohort.** First, ASVs identified in the validation cohort were matched to ASVs assigned to the 27 CAGs in the discovery cohort based on identical ASV sequences. Matched ASVs inherited the CAG assignments of their corresponding discovery cohort ASVs, and read counts were aggregated within each CAG. Relative abundances of the 27 CAGs were then calculated for each sample by dividing aggregated CAG counts by the total sequencing reads.

To assess whether the CAG framework was preserved following transfer, mean relative abundances of the 27 CAGs were compared between the discovery and validation cohorts. The resulting CAG profiles showed a high degree of concordance (Pearson's $r = 0.93$, $P < 0.001$), indicating that the overall CAG composition identified in the discovery cohort was broadly preserved in the validation cohort (Supplementary Figure S5).

To evaluate whether the functional characteristics associated with individual CAGs were preserved following transfer of the CAG framework, CAG–metabolite association coefficients obtained independently in the discovery and validation cohorts were compared. For each cohort, associations between the 27 CAGs and fecal metabolites were estimated by Spearman's rank correlation analysis using CAG relative abundances and metabolite concentrations. Correlation coefficients (ρ values) obtained for each CAG–metabolite pair were then compared between the two cohorts by Pearson correlation analysis. SCFAs and BAs were evaluated separately because bile acid profiles differed between the cohorts. Specifically, the validation cohort additionally included LCA-derived metabolites that were not measured in the discovery cohort. Therefore, only metabolites quantified in both cohorts were included in the cross-cohort comparison. Scatter plots comparing CAG–metabolite association coefficients between the D- and V-cohorts are shown in Supplementary Figure S6. Detailed CAG–metabolite association patterns in the V-cohort together with comparisons with the D-cohort are presented in Supplementary Figure S7.

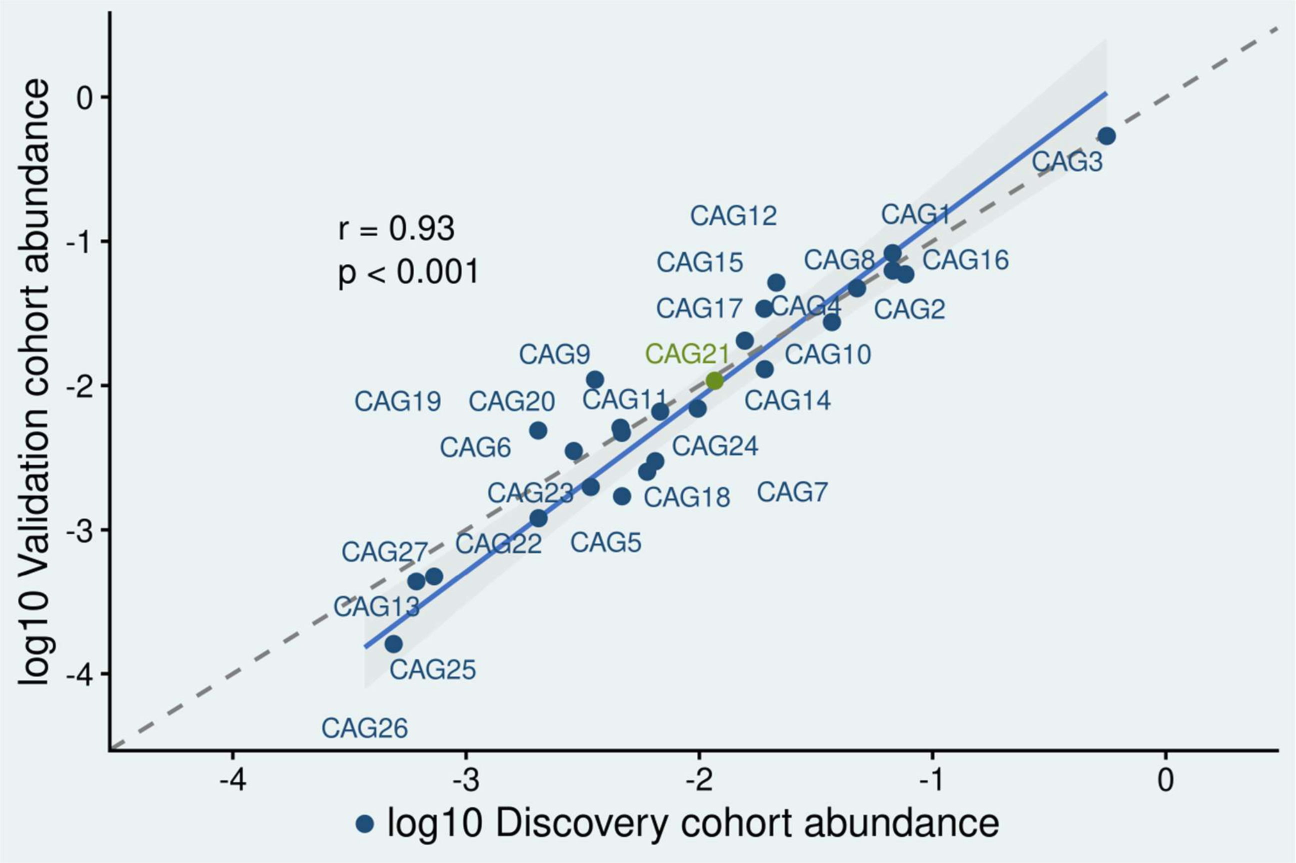


**Supplementary Figure S5. Concordance of cohort-level CAG abundance profiles between the D- and V-cohorts.** Scatter plot comparing the mean relative abundances of the 27 co-abundance groups (CAGs) between the discovery and validation cohorts. Mean relative abundances were calculated for each cohort and visualized on a log10 scale after addition of a pseudocount. Each point represents a single CAG and is labeled accordingly. The blue solid line indicates the fitted linear regression, the shaded area represents the 95% confidence interval, and the gray dashed line indicates the identity line ($y = x$). Pearson correlation showed strong concordance between cohorts ($r = 0.93$, $P < 0.001$).

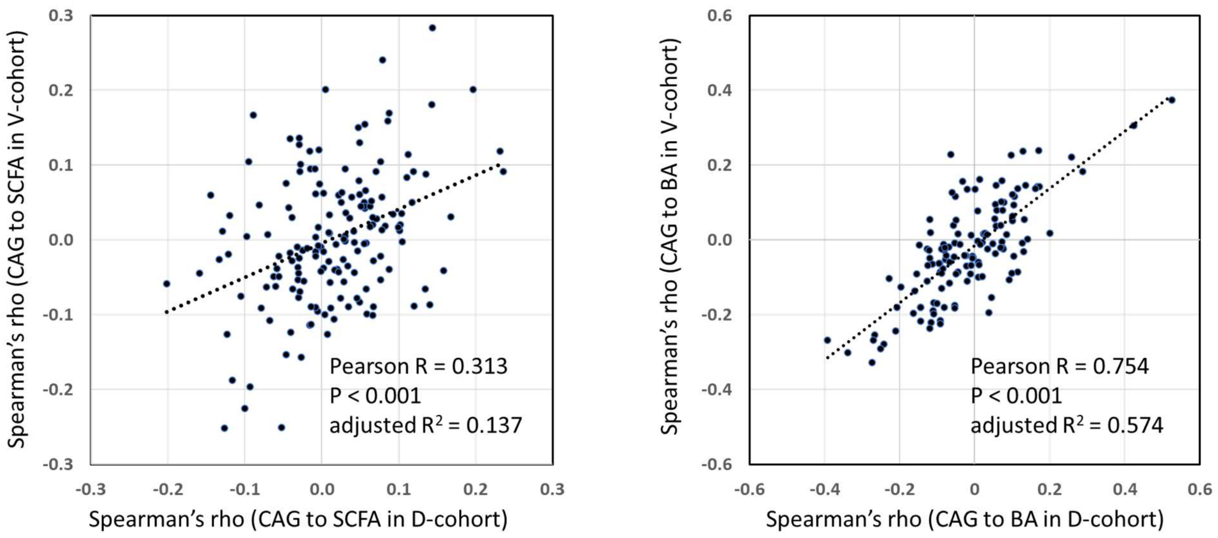


**Supplementary Figure S6.** Concordance of CAG–metabolite association profiles between the discovery and validation cohorts. Scatter plots comparing Spearman correlation coefficients for CAG–metabolite associations between the discovery and validation cohorts for (A) SCFAs and (B) BAs. Each point represents a single CAG–metabolite pair. Pearson correlation analysis demonstrated significant concordance of the overall association profiles for both SCFAs (R = 0.313, P < 0.001) and BAs (R = 0.754, P < 0.001).

| Module | acetate | propionate | butyarte | Succinate | Lactate | total SCFA | Conjugated BA | Primary BA (CA + CDCA) | Secondary BA (DCA + LCA) | UDCA | Tertiary BA | total BA |
|---|---|---|---|---|---|---|---|---|---|---|---|---|
| CAG1 | 0.012 | -0.004 | -0.046 | **0.101** | -0.068 | -0.014 | -0.052 | 0.078 | -0.050 | **0.162** | -0.064 | 0.013 |
| CAG2 | -0.093 | -0.027 | -0.038 | -0.052 | **-0.120** | **-0.100** | **-0.210** | **-0.267** | **0.288** | **-0.160** | **0.350** | -0.006 |
| CAG3 | **-0.121** | **-0.202** | **0.144** | -0.059 | 0.034 | **-0.133** | **-0.144** | **-0.250** | **0.167** | **-0.273** | **0.276** | -0.109 |
| CAG4 | 0.022 | -0.008 | 0.079 | -0.025 | -0.082 | 0.025 | -0.109 | **-0.145** | **0.135** | **-0.164** | **0.156** | -0.011 |
| CAG5 | **0.102** | 0.039 | **0.118** | 0.042 | 0.031 | 0.093 | -0.024 | -0.077 | 0.053 | **-0.113** | 0.069 | -0.010 |
| CAG6 | -0.043 | 0.003 | 0.077 | -0.003 | -0.008 | -0.029 | -0.082 | -0.089 | 0.070 | -0.044 | 0.091 | 0.015 |
| CAG7 | 0.049 | 0.046 | 0.087 | **-0.123** | -0.089 | 0.042 | **-0.242** | **-0.338** | **0.171** | **-0.392** | **0.317** | -0.096 |
| CAG8 | 0.019 | 0.030 | **-0.127** | **0.168** | **0.140** | 0.017 | **0.259** | **0.424** | **-0.269** | **0.526** | **-0.352** | **0.115** |
| CAG9 | **0.134** | **0.236** | 0.002 | **-0.129** | **-0.159** | **0.159** | -0.053 | **-0.127** | **0.129** | **-0.119** | 0.094 | 0.001 |
| CAG10 | 0.007 | -0.060 | **0.135** | 0.004 | 0.058 | -0.014 | 0.045 | **-0.120** | 0.097 | -0.110 | **0.168** | -0.027 |
| CAG11 | 0.029 | 0.027 | 0.058 | 0.063 | 0.002 | 0.055 | -0.004 | 0.014 | **-0.196** | -0.040 | **-0.217** | **-0.126** |
| CAG12 | 0.082 | **0.232** | 0.049 | -0.072 | **-0.145** | **0.119** | -0.066 | **-0.154** | **0.173** | **-0.207** | **0.157** | 0.027 |
| CAG13 | 0.056 | 0.030 | 0.086 | -0.015 | -0.023 | 0.049 | -0.069 | 0.020 | 0.057 | 0.091 | -0.004 | 0.048 |
| CAG14 | 0.010 | 0.031 | **0.120** | 0.077 | -0.028 | 0.022 | -0.033 | -0.063 | 0.099 | **-0.120** | **0.140** | 0.073 |
| CAG15 | 0.055 | **0.197** | -0.095 | 0.010 | -0.030 | 0.071 | -0.084 | **-0.118** | 0.100 | **-0.121** | **0.119** | 0.032 |
| CAG16 | 0.041 | **-0.116** | 0.016 | -0.015 | 0.005 | 0.009 | -0.020 | 0.100 | **-0.228** | 0.105 | **-0.211** | -0.076 |
| CAG17 | -0.062 | -0.070 | -0.056 | 0.066 | 0.068 | -0.079 | **0.140** | **0.126** | -0.067 | **0.200** | **-0.113** | 0.070 |
| CAG18 | 0.059 | 0.066 | -0.030 | -0.056 | -0.041 | 0.068 | 0.010 | 0.008 | 0.057 | 0.056 | 0.000 | 0.074 |
| CAG19 | -0.008 | 0.009 | -0.008 | -0.031 | -0.016 | -0.005 | -0.081 | -0.092 | 0.074 | -0.053 | 0.083 | 0.037 |
| CAG20 | -0.031 | -0.027 | -0.001 | 0.020 | -0.005 | -0.019 | 0.008 | -0.056 | 0.034 | -0.060 | 0.024 | -0.009 |
| CAG21 | **0.104** | **0.143** | -0.039 | **0.112** | 0.057 | **0.099** | **0.115** | **0.131** | -0.088 | 0.086 | -0.087 | -0.006 |
| CAG22 | 0.072 | **0.110** | **0.104** | -0.001 | -0.029 | 0.066 | -0.053 | -0.088 | 0.069 | -0.093 | 0.056 | 0.009 |
| CAG23 | 0.078 | 0.077 | 0.049 | 0.088 | -0.046 | 0.063 | -0.092 | 0.020 | 0.023 | -0.020 | 0.052 | 0.011 |
| CAG24 | 0.055 | 0.047 | 0.051 | -0.009 | 0.078 | 0.065 | 0.055 | 0.105 | 0.076 | 0.104 | -0.024 | **0.133** |
| CAG25 | 0.036 | 0.034 | -0.030 | 0.056 | 0.048 | 0.029 | 0.072 | 0.059 | -0.092 | 0.100 | -0.079 | -0.051 |
| CAG26 | -0.031 | -0.005 | -0.008 | -0.041 | -0.041 | -0.031 | **-0.116** | **-0.143** | -0.021 | **-0.147** | 0.052 | -0.100 |
| CAG27 | 0.024 | 0.028 | -0.097 | **-0.105** | -0.038 | 0.011 | -0.085 | -0.054 | -0.080 | -0.045 | -0.035 | **-0.125** |
| Correlation to D-cohort | | | | | | | | | | | | |
| R | 0.324 | **0.648** | **0.375** | 0.281 | -0.063 | **0.459** | **0.524** | **0.800** | **0.789** | **0.975** | ND | 0.439 |
| P | 0.076 | 0.000 | 0.001 | 0.092 | 0.671 | 0.007 | 0.000 | 0.000 | 0.000 | 0.000 | ND | 0.001 |

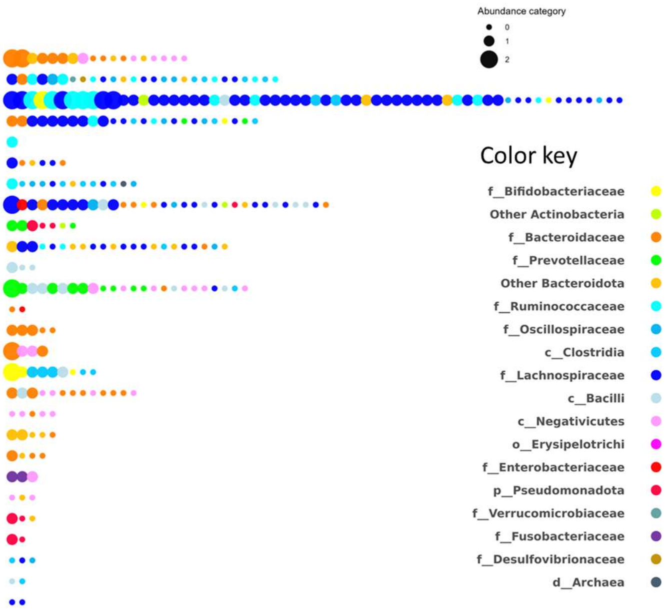


**Supplementary Figure S7.** CAG–metabolite associations in the validation cohort and taxonomic composition of individual CAGs. Left, heatmap showing Spearman's correlation coefficients between individual CAGs and each metabolite in the validation cohort. Positive and negative correlations are indicated by red and blue shading, respectively. Bold numbers indicate statistically significant associations ($P < 0.05$). The bottom row shows Pearson correlation coefficients comparing the CAG–metabolite association coefficients between the discovery and validation cohorts for each metabolite. Right, taxonomic composition of each CAG. Each dot represents an ASV assigned to a CAG, with dot size indicating its abundance category and dot color indicating its taxonomic affiliation at the family (or higher) level.

**2. Assignment of V-cohort samples to predefined GMTs.** To classify validation samples, cluster centroids representing each GMT were calculated from the discovery cohort. For each GMT, CLR-transformed relative abundance profiles of the 27 CAGs were averaged across all D-cohort samples assigned to that GMT (Supplementary Table S18). Prior to CLR transformation, a pseudocount of 0.5 was added to all CAG abundances to avoid zero values. The same CLR transformation was applied to validation cohort CAG profiles using an identical pseudocount. Euclidean distances between each validation sample and the seven discovery cohort centroids were calculated, and each sample was assigned to the GMT corresponding to the nearest centroid.

To assess assignment confidence, the Euclidean distance to the closest centroid, the distance to the second-closest centroid, and the margin between these two distances were calculated for every sample. The median margin between the nearest and second-nearest centroid distances was 0.07, with only 43 of 404 validation samples (10.6%) showing a margin of less than 0.01. These values were recorded but were not used to exclude any samples, ensuring that GMT classification in the validation cohort was performed without modifying either the CAG definitions or the GMT centroids established in the D-cohort.

**3. Evaluation of the transferred GMT configuration.** To evaluate the validity of the transferred GMT configuration, several complementary indices were calculated based on the assigned GMTs in the validation cohort.

First, the overall separation of the predefined GMTs was assessed using permutational multivariate analysis of variance (PERMANOVA) based on the Bray–Curtis distance matrix with 999 permutations. The proportion of variance explained by GMT identity ($R^2$) was used as a measure of the extent to which the transferred GMT configuration accounted for variation in microbial community composition (Supplementary Table S23).

Second, assignment confidence was evaluated from the centroid-based classification procedure. For each sample, Euclidean distances to the nearest and second-nearest GMT centroids were calculated in CLR-transformed CAG space, and the margin between these distances was recorded. The median assignment margin was 0.071, and only 43 of the 404 validation samples (10.6%) showed a margin of less than 0.01. These values were used only to evaluate assignment confidence and were not used to exclude any samples (Supplementary Table S24).

Finally, the compactness and separation of the transferred GMT configuration were assessed using silhouette analysis based on Euclidean distances in CLR-transformed CAG space. Silhouette coefficients were calculated for all validation samples, and the mean silhouette coefficient was used as an overall measure of clustering quality (Supplementary Table S25).

**Supplementary Table S23.** PERMANOVA analysis of GMT separation in the V-cohort.

| term | Df | SumOfSqs | R2 | F | Pr(>F) |
|---|---|---|---|---|---|
| Model | 7 | 41.58 | 0.454 | 74.73 | 0.001 |
| Residual | 628 | 49.92 | 0.546 | | |
| Total | 635 | 91.49 | 1.000 | | |

**Supplementary Table S24.** Assignment quality metrics for V-cohort GMT classification.

| assigned_cluster | n | best_distance_median | margin_median | frac_margin_lt_0_01 |
|---|---|---|---|---|
| MT-1 | 71 | 0.2635 | 0.0516 | 0.2113 |
| MT-2 | 37 | 0.2763 | 0.0467 | 0.0811 |
| MT-3 | 80 | 0.2025 | 0.0603 | 0.0750 |
| MT-4 | 36 | 0.3979 | 0.1087 | 0.0556 |
| MT-5 | 24 | 0.2309 | 0.0724 | 0.0417 |
| MT-6 | 90 | 0.1829 | 0.0576 | 0.1444 |
| MT-7 | 66 | 0.2388 | 0.1946 | 0.0455 |
| ALL | 404 | 0.2257 | 0.0710 | 0.1064 |

**Supplementary Table S25.** Silhouette index of GMTs in the validation cohort.

| GMT | n | mean_sil | median_sil | n | mean_sil | median_sil |
|---|---|---|---|---|---|---|
| MT-1 | 50 | -0.0336 | -0.0256 | 71 | 0.0203 | 0.0111 |
| MT-2 | 39 | -0.1055 | -0.1132 | 37 | -0.0236 | -0.0414 |
| MT-3 | 32 | 0.1158 | 0.1282 | 80 | 0.1720 | 0.1987 |
| MT-4 | 15 | 0.1292 | 0.1868 | 36 | -0.0321 | -0.0554 |
| MT-5 | 25 | 0.1513 | 0.1397 | 24 | 0.1243 | 0.1350 |
| MT-6 | 57 | 0.1226 | 0.1346 | 90 | 0.2913 | 0.3073 |
| MT-7 | 14 | 0.1176 | 0.1803 | 66 | 0.2305 | 0.2894 |
| ALL | 232 | 0.0529 | 0.0431 | 404 | 0.1425 | 0.1568 |

**4. Assessment of transferability and generalizability of the CAG-based GMT framework across cohorts.** To evaluate whether the gut microbiome types (GMTs) identified in the discovery cohort retained their characteristic microbial configurations in the validation cohort, the mean relative abundance profiles of the 27 co-abundance groups (CAGs) were compared between cohorts for each GMT.

For each cohort, the mean relative abundance of each CAG was calculated separately within each GMT and standardized by Z-score transformation across GMTs to facilitate comparison of relative enrichment patterns (Supplementary Figure S8). GMT-specific CAG configurations were visually compared between the discovery and validation cohorts. Statistical enrichment of individual CAGs within each GMT was evaluated independently in each cohort using one-versus-rest analysis with MaAsLin2 and Benjamini–Hochberg false discovery rate correction ($q \leq 0.10$), and significantly enriched CAGs are indicated by asterisks in Supplementary Figure S8.

To quantify the preservation of GMT-specific CAG configurations, Pearson correlation coefficients were calculated between the corresponding 27-dimensional CLR-transformed centroid vectors in the discovery and validation cohorts. Pearson correlation coefficients ranged from 0.63 to 0.81, indicating substantial preservation of GMT-specific CAG configurations across cohorts (Supplementary Table S26).

To evaluate whether the transferred GMT framework preserved the underlying CAG-based community structure across cohorts, samples from the discovery and validation cohorts were combined after assignment to the predefined GMTs. CLR-transformed CAG relative abundance profiles were used to calculate Euclidean distance matrices. Permutational multivariate analysis of variance (PERMANOVA) was performed using the adonis2 function in the vegan R package with 999 permutations. GMT assignment and cohort were included as explanatory variables. The proportion of explained variance ($R^2$) was used to compare the relative contributions of GMT and cohort to variation in CAG composition (Supplementary Table S27).

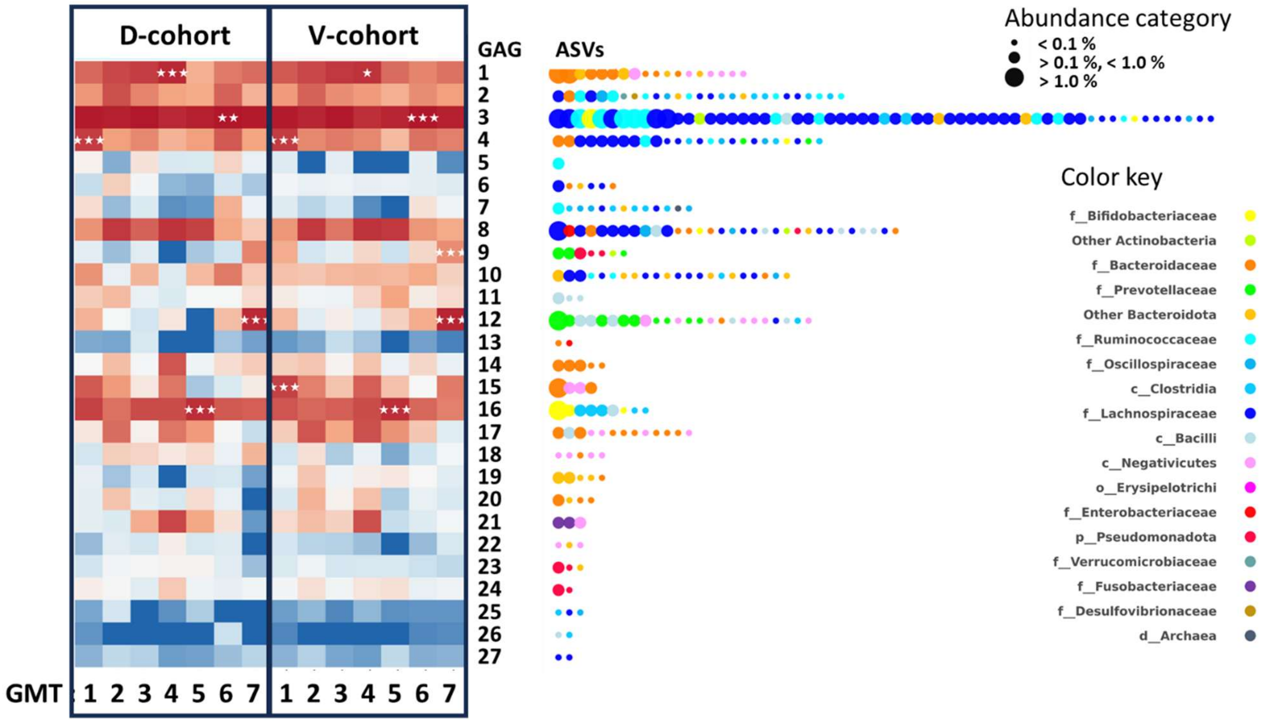


**Supplementary Figure S8. Preservation of GMT-specific CAG configurations between the discovery and validation cohorts.** Heatmaps showing the mean relative abundances of the 27 co-abundance groups (CAGs) across the seven gut microbiome types (GMTs) in the discovery cohort (left) and validation cohort (center). Relative abundances were standardized for each CAG using Z-score transformation to facilitate comparison across GMTs. Red and blue indicate relatively higher and lower abundances, respectively. White asterisks denote the GMT(s) in which individual CAGs were significantly enriched based on one-versus-rest analysis using MaAsLin2 with Benjamini–Hochberg false discovery rate correction ($q \leq 0.10$). The overall similarity of the GMT-specific CAG configurations between cohorts is summarized in Supplementary Table S27. The right panel shows the taxonomic composition of each CAG. Each dot represents a single ASV assigned to the corresponding CAG, with dot size indicating its mean relative abundance across the discovery cohort (<0.1%, 0.1–1.0%, or >1.0%) and dot color indicating its taxonomic affiliation at the family level or the lowest available higher taxonomic rank when family-level assignment was unavailable.

**Supplementary Table S26. Similarity of GMT-specific 27-CAG centroid profiles between D- and V-cohorts.**

| GMT | Pearson's r | Euclidean distance |
|---|---|---|
| MT-1 | 0.657 | 18.826 |
| MT-2 | 0.730 | 18.955 |
| MT-3 | 0.654 | 19.790 |
| MT-4 | 0.809 | 18.988 |
| MT-5 | 0.732 | 19.799 |
| MT-6 | 0.628 | 18.578 |
| MT-7 | 0.663 | 19.371 |

Pearson's correlation coefficients were calculated between the corresponding 27-dimensional CLR-transformed GMT centroid vectors in the discovery and validation cohorts. Euclidean distances were calculated in the same CLR-transformed space used for validation sample assignment. Higher Pearson's r and lower Euclidean distance indicate greater similarity between corresponding GMTs.

**Supplementary Table S27.** PERMANOVA of GMT and cohort effects on CAG composition in the combined discovery and validation cohorts.

| | Df | SumOfSqs | R2 | F | Pr(>F) |
|---|---|---|---|---|---|
| cluster | 6 | 41.234 | 0.4507 | 86.461 | 0.001 |
| cohort | 1 | 0.343 | 0.0037 | 4.310 | 0.001 |
| Residual | 628 | 49.917 | 0.5456 | | |
| Total | 635 | 91.494 | 1.0000 | | |

PERMANOVA was performed using Euclidean distances calculated from CLR-transformed relative abundance profiles of the 27 co-abundance groups (CAGs). The analysis included both GMT identity and cohort (discovery vs. validation) as explanatory variables.  P-values were calculated using permutation tests with 999 permutations.

##### 5. Assessment of cross-cohort reproducibility of GMT-associated metabolite profiles.

To evaluate whether GMT-associated metabolite profiles were preserved between the discovery and validation cohorts, mean metabolite profiles were calculated separately for each GMT in each cohort. For bile acids, relative abundances of metabolite groups were used, whereas absolute concentrations were used for SCFAs. Bray–Curtis dissimilarity matrices were calculated among the seven GMTs within each cohort using the vegan package in R. Concordance between the discovery- and validation-cohort distance matrices was assessed using Spearman Mantel tests with up to 9,999 permutations. Because only seven GMTs were compared, the number of unique permutations was limited, and all possible permutations were evaluated where applicable.

#### Supplementary Note 9. Association analyses between GMTs and host clinical characteristics.

**1. Clinical parameters.** Associations between GMTs and host clinical characteristics were evaluated using a predefined panel of 11 clinical parameters comprising BMI, HbA1c, triglycerides (TG), HDL cholesterol, LDL cholesterol, γ-glutamyl transpeptidase (γ-GTP), alanine aminotransferase (ALT), estimated glomerular filtration rate (eGFR), uric acid (UA), systolic blood pressure (SBP), and diastolic blood pressure (DBP). TG and γ-GTP values were natural log-transformed after addition of a small pseudocount, and all clinical variables were standardized to zero mean and unit variance (z-score transformation). Samples with missing values in any clinical parameter were excluded by complete-case analysis.

**2. Distance-based redundancy analysis.** For multivariate analysis of host clinical profiles, Euclidean distances were calculated from the standardized clinical parameter matrix. Distance-based redundancy analysis (dbRDA) was performed using the capscale function in the vegan R package. GMT assignment was included as the primary explanatory variable, whereas age, sex, and medication status were included as additional explanatory variables. The significance of the overall model, individual explanatory variables, and constrained axes was assessed using permutation tests (999 permutations). Model explanatory power was summarized using $R^2$ and adjusted $R^2$. The resulting ordination plot is shown in Supplementary Figure S9.

**3. Multinomial logistic regression.** To evaluate whether associations between GMTs and host clinical characteristics were independent of demographic factors and medication use, multinomial logistic regression models were constructed using GMT assignment as the outcome variable. Three models were fitted: (i) a null model containing only the intercept, (ii) an adjustment model including age, sex, and medication status, and (iii) a full model including the clinical parameter panel in addition to the adjustment variables. Model fit was evaluated using McFadden's pseudo-$R^2$ and Nagelkerke's $R^2$. The incremental contribution of the clinical parameter panel beyond the adjustment model was assessed using likelihood ratio tests. Predictive performance of the full model was evaluated by five-fold cross-validation. The results of the multinomial logistic regression analysis are summarized in Supplementary Table S28.

**4. Estimation of MT-specific adjusted probabilities for individual clinical parameters**

To characterize MT-specific patterns in individual clinical parameters, binary logistic regression analyses were performed using Stata/SE 12 (StataCorp, College Station, TX, USA). Each dichotomized clinical parameter was analyzed separately as the dependent variable, with GMT assignment included as a categorical explanatory variable and age included as a covariate. Following model fitting, adjusted probabilities of meeting the criterion for each clinical outcome were estimated for individual GMTs using the postestimation margins command. The overall prevalence of each clinical outcome was calculated across the analyzed subjects and used as a reference for comparison with the GMT-specific adjusted probabilities. Adjusted probabilities and their 95% confidence intervals were plotted for each GMT, with the overall prevalence of the corresponding clinical outcome shown as a reference line.

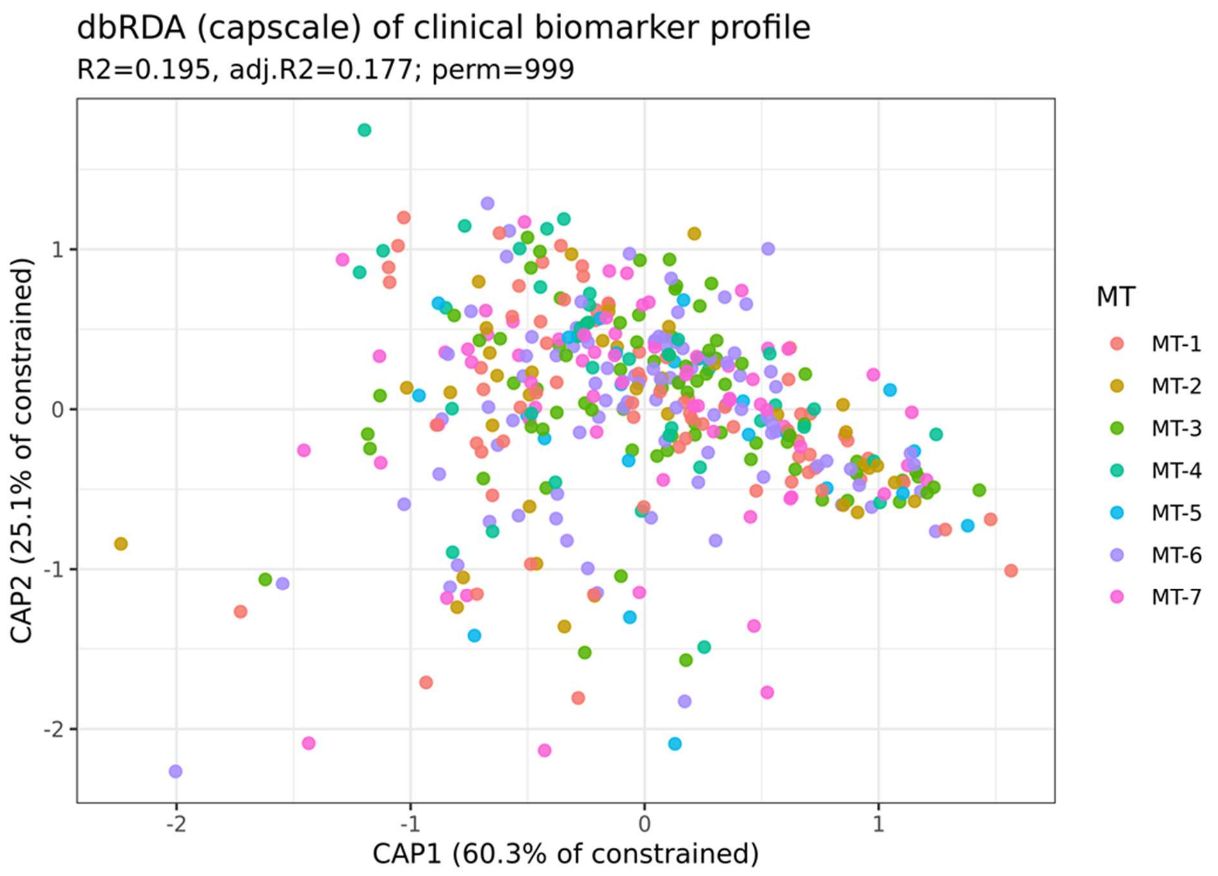


**Supplementary Figure S9. Distance-based redundancy analysis (dbRDA) of clinical biomarker profiles.** dbRDA was performed using the predefined clinical biomarker panel, including anthropometric, glycemic, lipid, liver, renal, and blood pressure parameters. Samples are colored according to GMT assignment. The constrained ordination model explained 19.5% of the total variation (adjusted $R^2$ = 17.7%). Although GMTs showed considerable overlap in the ordination space, permutation testing identified a significant association between GMT assignment and multivariate clinical biomarker profiles.

| **Supplementary Table S28.** Comparison of multinomial logistic regression model performance for the association between clinical parameters and microbiome-type assignment. | | | | | |
|---|---|---|---|---|---|
| **Clustering**[a] | **Predictors**[b] | **Adjustment** | **McFadden's pseudo-R² c** | **Nagelkerke's R²** | **LR test P**[d] |
| 7 GMTs / 27 CAGs | — | age, sex, medication | 0.023 | 0.085 | — |
| 7 GMTs / 27 CAGs | Clinical panel[c] | age, sex, medication | 0.092 | 0.298 | — |
| 7 GMTs / 27 CAGs | Clinical panel | age, sex, medication | Δ 0.070 | — | 0.002 |
| 7 GMTs / 27 CAGs | Clinical panel | medication | 0.081 | 0.266 | 0.000 |
| 7 GMTs / 229 genera | Clinical panel | age, sex, medication | 0.097 | 0.296 | 0.018 |

| **Additional validation** | **Value** |
|---|---|
| 5-fold CV accuracy | 0.205 |
| Balanced accuracy | 0.161 |

[a]7 GMTs/27 CAGs: 7GMTs clustered based on 27 CAGs transferred from discovery cohort; 7 GMTs/229 genera: 7 GMTs clustered based on 229 genera with prevalence > 1% in the validation cohort.

[b]Clinical panel: BMI, HbA1c, TG(log), HDL, LDL, γGTP(log), ALT, eGFR, UA, SBP, and DBP.

[c]Δ indicates the incremental increase in McFadden's pseudo-$R^2$ after addition of the clinical panel to the adjustment model.

[d]LR test P indicates the P-value from the likelihood ratio test.

### Supplementary Note 10. Comparative evaluation of the CAG-based GMT framework with a genus-based framework

**1. Comparison with de novo genus-based microbiome stratification.** To compare the performance of the transferred CAG-based GMT framework with a conventional taxonomic stratification approach, an independent genus-based microbiome classification was constructed de novo using the validation cohort. Genus-level relative abundance profiles were used as input for microbiome clustering, and each validation cohort sample was assigned to a genus-based microbiome type according to the resulting clustering. The resulting genus-based microbiome types were evaluated using the same multinomial logistic regression framework applied to the transferred CAG-based GMT classification. Model performance was assessed using likelihood ratio tests, McFadden's pseudo-$R^2$, Nagelkerke's $R^2$, and five-fold cross-validation, allowing direct comparison between the transferred CAG-based framework and the independently derived genus-based classification. The comparative performance metrics are summarized in Supplementary Table S28.

**2. Comparison of explanatory value between GMTs and representative genera,** To evaluate whether host-associated clinical phenotypes were better explained by microbiome community states than by individual bacterial taxa, additional Firth logistic regression analyses were performed for selected GMT–clinical associations. MT-4 and MT-5 were selected because they exhibited the strongest and most reproducible associations with liver and renal function, respectively. *Faecalibacterium* and *Bifidobacterium*, which consistently characterized MT-4 and MT-5 in both the discovery and validation cohorts, were selected as representative genera.

For each GMT–clinical association, three Firth logistic regression models were fitted after adjustment for age, sex, and medication use: (i) a GMT-only model, (ii) a representative genus-only model, and (iii) a combined model including both the GMT and its corresponding representative genus. Nested likelihood ratio tests were performed to determine whether inclusion of either the GMT or representative genus significantly improved model fit beyond the corresponding comparison model. For the MT-4 analysis, alcohol consumption was additionally included as a covariate because of its significant association with both MT-4 and serum γ-glutamyl transpeptidase (γ-GTP).

**2. Comparison of explanatory value between GMTs and representative genera.** To evaluate whether host-associated clinical phenotypes were better explained by microbiome community states than by individual bacterial taxa, additional Firth logistic regression analyses were performed for selected GMT–clinical associations. MT-4 and MT-5 were selected because they exhibited the strongest and most reproducible associations with liver and renal function, respectively. *Faecalibacterium* and Bifidobacterium, which consistently characterized MT-4 and MT-5 in both the discovery and validation cohorts, were selected as representative genera.

For each GMT–clinical association, three Firth logistic regression models were fitted after adjustment for age, sex, and medication use: (i) a GMT-only model, (ii) a representative genus-only model, and (iii) a combined model including both the GMT and its corresponding representative genus. The results of these analyses are summarized in Supplementary Table S29.

Nested likelihood ratio tests were performed to determine whether inclusion of either the GMT or representative genus significantly improved model fit beyond the corresponding comparison model. For the MT-4 analysis, alcohol consumption was additionally included as a covariate because of its significant association with both MT-4 and serum γ-glutamyl transpeptidase (γ-GTP). The results of the nested likelihood ratio tests are summarized in Supplementary Table S30.

**Supplementary Table S29.** Representative genus-adjusted Firth logistic regression analyses of GMT–clinical associations.

| GMT | Clinical parameter | Model | MT OR (95% CI) | MT P | Genus OR (95% CI) | Genus P |
|---|---|---|---|---|---|---|
| MT-4 | γGTP | MT-4 | 2.33 (1.15–4.77) | **0.019** | — | — |
| MT-4 | γGTP | *Faecalibacterium* | — | — | 0.86 (0.70–1.06) | 0.153 |
| MT-4 | γGTP | MT-4 + *Faecalibacterium* | 2.14 (0.99–4.69) | 0.055 | 0.94 (0.75–1.20) | 0.614 |
| MT-5 | eGFR | MT-5 | 0.085 (0.00067-0.64) | **0.0099** | - | - |
| MT-5 | eGFR | *Bifidobacterium* | — | — | 0.78 (0.62-0.92) | 0.035 |
| MT-5 | eGFR | MT-5 + *Bifidobacterium* | 0.104 (0.00081-0.80) | **0.024** | 0.82 (0.65-1.04) | 0.101 |

Binary outcomes were defined using the predefined clinical cutoffs described in the Methods. Firth logistic regression models were fitted with age, sex, and medication use as covariates. For each GMT–clinical association, three models were evaluated: (i) GMT only, (ii) the genus characterizing the corresponding GMT only, and (iii) GMT plus the genus characterizing the corresponding GMT. Representative genera were defined as genera consistently characterizing the corresponding GMTs in both the discovery and validation cohorts.

**Supplementary Table S30.** Nested likelihood ratio tests evaluating the relative explanatory contributions of MT-4 and Faecalibacterium abundance to elevated γ-glutamyl transpeptidase (γ-GTP). Logistic regression models were adjusted for age, sex, medication use, and alcohol consumption. Likelihood ratio $\chi^2$ statistics represent the reduction in deviance obtained by adding the indicated variable to the corresponding baseline model.

| Comparison | LR$\chi^2$ | P value |
|---|---|---|
| Faecalibacterium added to the covariate model | 1.574 | 0.21 |
| MT-4 added to the covariate model | 5.874 | 0.015 |
| Faecalibacterium added to the MT-4 model | 0.043 | 0.836 |
| MT-4 added to the Faecalibacterium model | 4.342 | 0.037 |
